# Post-translational Tuning of Human Cortical Progenitor Neuronal Output

**DOI:** 10.1101/2025.10.27.684791

**Authors:** Julien Pigeon, Tamina Dietl, Myriame Abou Mrad, Ludovico Rizzuti, Miguel V. Silva, Natasha Danda, Corentine Marie, Clarisse Brunet Avalos, Hayat Mokrani, Laila El Khattabi, Alexandre D. Baffet, Diogo S. Castro, Carlos Parras, Boyan Bonev, Bassem A. Hassan

## Abstract

The striking expansion of the human neocortex is driven by the ability of radial glial cells (RGCs) to prolong neurogenesis without exhausting their progenitor pool, thereby producing more neurons, particularly in the upper layers. The mechanisms underlying enhanced human neurogenic output remain unresolved. We hypothesized that post-translational modification of conserved proteins could provide a flexible mechanism supporting species-specific patterns of neurogenesis. Here, we identify phosphorylation of the proneural factor NEUROGENIN2 as a regulator of human RGC neuronal output by modulating its pioneering-like activity, at the expense of its transactivation function. This, in turn, remodels the RGC chromatin landscape, resulting in a forward temporal shift of RGC gene regulatory networks and leading to increased neuronal production biased towards upper-layer neurons. Our work proposes a conceptual framework for how subtle modifications of conserved processes, and not just new genes, encode tunable mechanisms for species-specific trajectories of brain development.

## INTRODUCTION

The evolutionary expansion of the human neocortex underlies higher-order cognitive abilities, such as reasoning, language, and abstract thought ^1,2^. Architecturally defined by six layers of neurons formed in an inside-out sequence, with early generation of deep layer neurons, followed by later generation of upper layer neurons^3–5^, the human neocortex contains more neurons than any other species, with approximately 16 billion in humans compared to 6 billion in chimpanzees, 1.7 billion in rhesus macaques, 31 million in rats, and 14 million in mice ^6–8^. This expansion of the primate neocortex is most pronounced in the upper layers, which show increases in both neuron number and molecular subtype diversity ^9–12^. Increased neurogenesis is facilitated by a prolonged neurogenic period^13–16^ during which cortical progenitors called Radial Glial Cells (RGCs) engage in a temporally controlled balance between proliferative divisions that expand their pool and neurogenic divisions that generate neurons either directly from RGCs (direct neurogenesis), or indirectly through the generation of intermediate progenitors (IPs; indirect neurogenesis). It remains unclear how human RGCs evolved their species-specific extension of the neurogenic period to increase neuronal output. Recent work has focused on human-specific evolutionary driving forces^17^ such as gene duplications of ARHGAP11B ^18–20^, NOTCH2NL ^21,22^, CROCCP2 ^23^, and SRGAP2C ^24,25^, and regulatory innovations with human-accelerated regions (HARs) like HAR5 ^26,27^. However, most of these genomic-level human-specific innovations do not generate new mechanisms. Instead, they converge on classic developmental pathways such as Notch, WNT, or PI3K-Akt ^28^, to modify the activity, timing, or expression of neurogenic programs that are otherwise shared across vertebrates. This raises a fundamental question: can human-specific features of cortical development arise from regulatory tuning of deeply conserved mechanisms, without changes to gene sequence? In particular, can we fine-tune the neuronal production of human cortical progenitors through post-slational modifications of conserved transcription factors that gate neurogenic programs?

In the developing mouse neocortex, Neurogenin2 (Neurog2), a highly conserved basic-helix-loop-helix (bHLH) transcription factor, is both necessary and sufficient for deep-layer cortical neurogenesis, but is dispensable for the generation of upper layer neurons ^29–34^, the main target of human cortical expansion. By integrating upstream signals such as Notch, FGF, and WNT into transcriptional programs, Neurog2 acts as a hub to promote neuronal differentiation and fate commitment during mouse neocortical development ^29,35–38^. Surprisingly, whether NEUROG2 is required for human cortical neurogenesis has not been investigated.

Proneural proteins, like NEUROG2, do not function as binary switches. Instead, their activity is finely tunable through post-translational modifications (PTMs) such as phosphorylation to modulate their stability, activity, and transcriptional output ^37,39–41^. A conserved phosphorylation site in NEUROG2, threonine 149 (T149), located at the junction between the loop and the second helix of its bHLH domain, directly modulates its neurogenic activity. Overexpression of a phospho-mutant form (T149A) of Neurog2 in developing mouse cortical RGCs enhances its neurogenic activity, promoting early neurogenesis and favoring deep-layer neuron production at the expense of upper-layer neurons ^39^. Given the species-specific expansion of human RGCs and their increased neuronal output, particularly in upper layers, we asked whether there may be a species-specific alteration to NEUROG2 function such that phosphorylation enables context-specific functional plasticity to modulate the neurogenic capacities of human RGCs. This would provide a tunable cell biological mechanism to extend neurogenic trajectories throughout neocortical development without requiring human-specific genetic changes.

To test this hypothesis, we used CRISPR/Cas9 genome editing in two different human iPSC lines to generate two different mutants: NEUROG2 KO to ask whether NEUROG2 is required for human cortical neurogenesis, and NEUROG2 T149A to ask whether its function can be tuned by PTM. We then combined live single progenitor lineage tracing of ∼400 RGCs, and developed a machine learning based pipeline to quantify ∼100 million cells across ∼200 hCOs to characterize the division patterns and neurogenic output of control and NEUROG2 mutant human cortical RGCs. Next, we leveraged paired single-nucleus RNA-seq and ATAC-seq, Gene Regulatory Network (GRN) analysis, and experimental validation to discover the molecular mechanisms underpinning NEUROG2 T149 modulation in human RGCs. At the cellular level, we find that, in contrast to mice, NEUROG2 expression is largely restricted to RGCs and is required for both deep and upper layer neurogenesis. The T149 phosphorylation site acts as a rheostat of RGC division patterns to fine-tune the balance between proliferative and neurogenic divisions. At the molecular level, inhibiting T149 phosphorylation uncouples NEUROG2’s chromatin modulating activity from its transcriptional activity, promoting the opening of different binding site of the AP-1 complex (JUN and FOS) specifically in RGCs. This alters RGC gene regulatory networks, driving enhanced neurogenic commitment and leading to increased production of upper-layer neurons. Together, our results support the hypothesis that conserved and tunable cell biological mechanisms can underlie species-specific expansion in brain size.

## RESULTS

### Human NEUROG2 is largely restricted to RGCs

To investigate neurogenesis in hCOs, we used a 3D cortical differentiation protocol (Figure S1A) adapted from Sloan et al.,^42^ and we confirmed the presence of all key cell subtypes found in the developing human neocortex (Figure S1C-G). At day 70, we found all progenitor subtypes, such as radial glia cells (RGCs) expressing SOX2 and PAX6 (Figure S1C & S1D), and Intermediate progenitors (IPs), expressing TBR2 (Figure S1C & S1D). At this stage of cortical development, these progenitors mostly produce TBR1+ deep-layer neurons and are just starting to produce SATB2+ upper-layer neurons (Figure S1E). However, at later stages of neocortical development, around day 140, these SOX2+ progenitors produce both deep and upper-layer neurons (Figure S1F & S1G). The high number and high density of cells in 3D hCOs pose a significant challenge for quantification; thus, we developed a semi-automated image analysis pipeline integrating Deep Learning (DL) and Machine Learning (ML) algorithms to measure cell proportions with precision (Figure 1a, see Methods for details). We fine-tuned the StarDist segmentation model ^43,44^ to better detect individual nuclei in very dense regions of the hCOs where we found that the F1 score improved from 0.64 to 0.83, indicating a substantial increase in segmentation accuracy (Figure S2B-E). Next, we used an ML classifier based on random forests in the open-source software *ilastik*^45^ to categorize each nucleus into distinct subtypes defined by standard cell-type markers. This image analysis pipeline allowed us to quantify thousands of sections containing hundreds of thousands of cells. In all the experiments described below, all nuclei within each section of each organoid used in this study were quantified to obtain the data presented.

**Figure 1.**
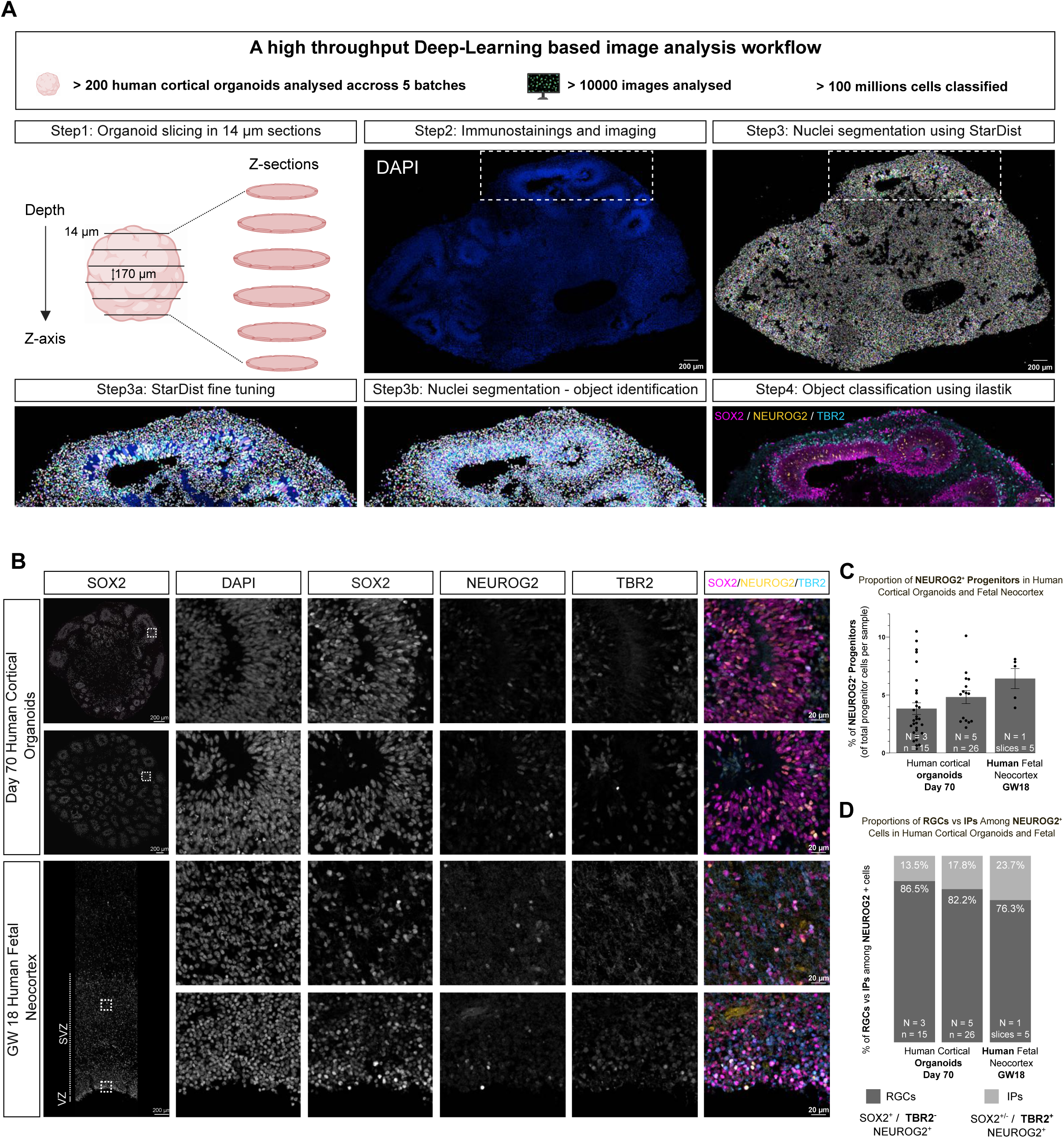
High-throughput deep learning-based image analysis of cell subtypes in human cortical organoids and fetal neocortex. **(A)** Overview of the high-throughput image analysis workflow. Step 1: Organoids were sectioned at 14 μm thickness to generate multiple slices across the tissue depth. Step 2: Sections were immunostained and imaged for DAPI and relevant markers. Step 3: Nuclei were segmented using StarDist, a deep learning-based algorithm. Step 3a: initial segmentation using the pretrained StarDist model. Step 3b: segmentation refined using a custom fine-tuned StarDist model (haug2). Step 4: Cells were classified based on marker expression using ilastik. The pipeline was applied to 200 organoids across 5 independent differentiations, analyzing >10,000 images and classifying over 100 million cells. **(B)** Representative confocal images of two wild-type clones of iPSC-derived cortical organoids at day 70 (top rows) and GW18 human fetal neocortex (bottom rows, corresponding to the subventricular and ventricular zones), stained for DAPI, SOX2, NEUROG2, and TBR2. Merged images show SOX2 (magenta), NEUROG2 (yellow), and TBR2 (cyan). Dotted boxes indicate regions shown at higher magnification on the right. Scale bars, 200 μm (overview), 20 μm (zoom). **(C)** Quantification of NEUROG2+ progenitors, performed using the pipeline described in (A), expressed as a percentage of total SOX2+ and/or TBR2+ progenitor cells in day 70 human cortical organoids and GW18 fetal neocortex. **(D)** Proportion of radial glia cells (RGCs, SOX2+TBR2−) and intermediate progenitors (IPs, TBR2+ and SOX2+/-) among all NEUROG2+ cells in human cortical organoids and fetal neocortex. N = number of biological replicates (organoid differentiations or embryos); n/slices = number of organoids or tissue sections analyzed.

We next examined the expression pattern of NEUROG2 in the progenitor populations of hCOs at day 70 through immunostaining for NEUROG2, SOX2 (pan-progenitor marker, RGCs and IPs), and TBR2 (intermediate progenitor marker, IPs only) (Figure 1B). In two independent clones, we observed that 4-5% of all progenitors expressed NEUROG2 (Figure 1C). To determine the identity of the cells expressing NEUROG2, we quantified the proportion of RGCs (SOX2+/TBR2-) and IPs (TBR2+) within the NEUROG2+ population. In both sets of control organoids, the majority of NEUROG2+ cells were RGCs (86.5% and 82.2%), while a smaller fraction corresponded to IPs (13.5% and 17.8%) (Figure 1D). To assess the relevance of these findings to human brain development, we next examined NEUROG2 expression in the Human Fetal Neocortex (HFC). We found similar proportions in the developing HFC at gestational week 18, developmentally comparable to D70-90 in hCOs, where 6% of all progenitors expressed NEUROG2 (Figure 1C). Among these NEUROG2+ cells, 76.3% were RGCs and 23.7% were IPs (Figure 1D).

Together, these results indicate that in the human developing neocortex, NEUROG2 is predominantly expressed in RGCs, a notable divergence from the murine developing neocortex, where NEUROG2 expression predominates in IPs at mid-stage of neocortical development^34,46^.

### NEUROG2 is required for efficient and timely human cortical neurogenesis

Given this species-specific difference, we sought to examine the role of ENUROG2 in human cortical neurogenesis. In the developing mouse neocortex, NEUROG2 is considered the main driver of deep-layer cortical neurogenesis, but it is dispensable for the generation of upper-layer neurons ^34,46^. To this end, we used CRISPR/Cas9 genome editing to generate from our main iPSC line, WTSli008-A, two independent NEUROG2 knockout iPSC clones referred to as KO1 and KO2, each carrying early stop codons (Figure S3A). A third clone (Ctrl), exposed to the editing protocol but unmodified, served as the isogenic control. We validated the NEUROG2 KO in a different genetic background with the GM25256*E iPSCs line (Figure S3D). We first validated in the main iPSC line (WTSli008-A) the loss of NEUROG2 protein expression and assessed early progenitor dynamics at D70 of differentiation by immunostaining COs for SOX2 (pan-progenitor marker), TBR2 (IP marker), and NEUROG2 (Figure 2A). In control organoids, 4.9% ± 0.5% of progenitors expressed NEUROG2, whereas NEUROG2 expression was reduced to background levels in KO1 (0.61% ± 0.5%) and KO2 (0.26% ± 0.5%) organoids (Figure S3B & S3C), confirming the loss of NEUROG2 in these clones.

**Figure 2:**
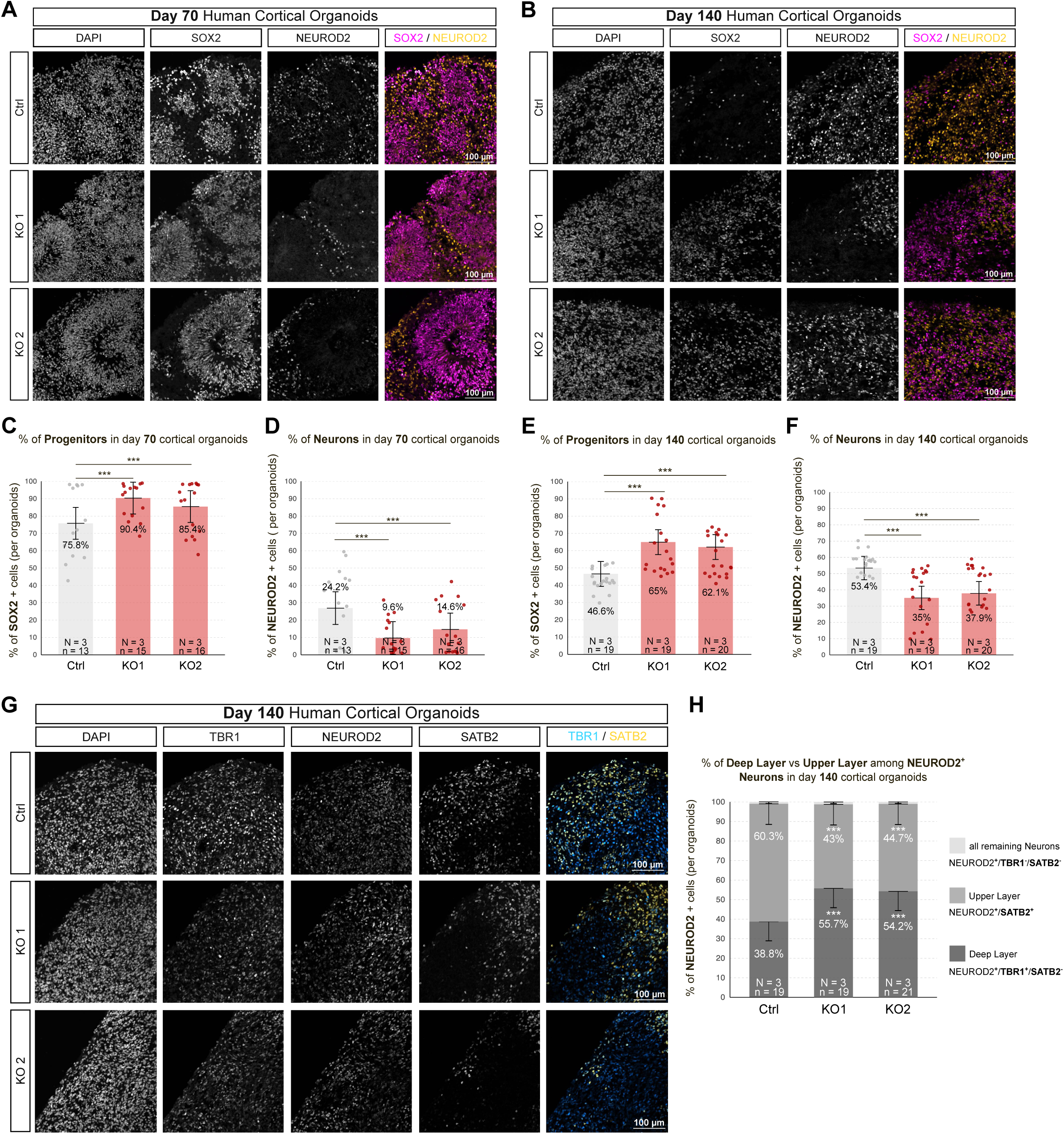
NEUROG2 regulates human cortical neurogenesis. **(A–B)** Representative confocal images of human cortical organoids at day 70 **(A)** and day 140 **(B)**, stained for SOX2 (magenta), NEUROD2 (yellow), and DAPI (gray). Ctrl: wild-type clone; KO1 and KO2: NEUROG2 knockout clones. Scale bars, 100 μm. **(C)** Quantification of the proportion of SOX2⁺ progenitors at day 70, expressed as a percentage of all antibody-stained cells. **(D)** Quantification of the proportion of NEUROD2⁺ neurons at day 70, expressed as a percentage of all antibody-stained cells. **(E)** Quantification of the proportion of SOX2⁺ progenitors at day 140, expressed as a percentage of all antibody-stained cells. **(F)** Quantification of the proportion of NEUROD2⁺ neurons at day 140, expressed as a percentage of all antibody-stained cells. **(G)** Representative confocal images of day 140 organoids stained for TBR1 (cyan), SATB2 (yellow), and NEUROD2 (gray). Scale bars, 100 μm. **(H)** Quantification of the proportion of deep-layer (TBR1⁺) and upper-layer (SATB2⁺) neurons among all NEUROD2⁺ cells, expressed per organoid.Bar graphs show adjusted means ± SEM derived from a linear mixed-effects model (group as fixed effect; differentiation batch as random effect). Pairwise comparisons were performed using estimated marginal means (emmeans); outliers (residuals > 2 SD) were excluded. N = number of independent differentiations; n = number of organoids. p < 0.05; p < 0.01; p < 0.001.

To assess the consequences of NEUROG2 depletion on neurogenesis, we quantified the proportions of progenitors and neurons in hCOs at mid (D70) and late (D140) stages using immunostaining for SOX2 (progenitors) and NEUROD2 (excitatory neurons) (Figure 2A & 2B). At mid stages, in control organoids we found that 75.8% ± 9.2% of the cells were progenitors and 24.2% ± 9.2% were neurons (Figure 2c, d). In contrast, KO clones showed a significant increase in progenitors (KO1 90.4% ± 9.2%, KO2 85.4% ± 9.2%) and an approximately 2-fold reduction in neurons (KO1 9.6% ± 9.2%, KO2 14.6% ± 9.2%) (Figure 2C, D). These results were also validated in our second iPSC cell line (Figure S3E-G). At later stages, control organoids contained 46.6% ± 7.2% progenitors and 53.4% ± 7.2% neurons, whereas KO organoids displayed a persistent neurogenic deficit, with significantly elevated proportions of progenitors (KO1: 65.0% ± 7.2%; KO2: 62.1% ± 7.2%) and a ∼1.5 fold reduction in neurons (KO1 35.0% ± 7.2%, KO2 37.9% ± 7.2%) (Figure 2E, F). These findings indicate a sustained impairment in neurogenic output and prolonged maintenance of progenitor identity upon NEUROG2 loss in hCOs.

To examine whether the loss of NEUROG2 specifically affects the generation of deep-layer neurons, as previously found in mice^32^, we quantified neuronal subtype identity among all NEUROD2+ cells using TBR1 (deep layer) and SATB2 (upper layer) at D140 (Figure 2G). In control organoids, 60.3% ± 10.6% neurons were SATB2[upper-layer neurons, while the remaining 38.8% ± 9.8% were TBR1[deep-layer neurons (Figure 2H). In contrast, KO1 and KO2 organoids exhibited a shift toward deep-layer identity, with an increased proportion of TBR1[neurons (KO1: 55.7% ± 9.8%; KO2: 54.2% ± 9.8%) and a corresponding reduction in upper-layer neurons (KO1: 43.0% ± 10.5%; KO2: 44.7% ± 10.5%) (Figure 2H).

Thus, in contrast to the mouse, human NEUROG2 is required for the generation upper-layer neurons.

### A conserved phosphorylation site regulates NEUROG2 proneural potency

Given the critical role of NEUROG2 in human cortical neurogenesis, we investigated mechanisms controlling its activity. Previous work in mice showed that phosphorylation of a conserved serine/threonine residue within the bHLH domain, the threonine 149 (T149) in NEUROG2, acts as a regulatory switch for the transcriptional and proneural potency of proneural proteins ^39^. This residue is conserved from insects to primates, including humans (Figure 3A), suggesting a strong evolutionary pressure to maintain its function. The gnomAD database comprising approximately 1.6 million sequenced alleles, identifying that NEUROGENIN2 T149 is rarely mutated. Only two heterozygous cases of a T149A substitution were detected where NEUROG2 cannot be phosphorylated anymore (Figure 3B). Moreover, pathogenicity predictions using AlphaMissense indicate that substitutions at this position are highly deleterious (score = 0.99, Figure 3B), among the most highly pathogenic in the bHLH domain highlighting its potential importance in human brain development.

**Figure 3:**
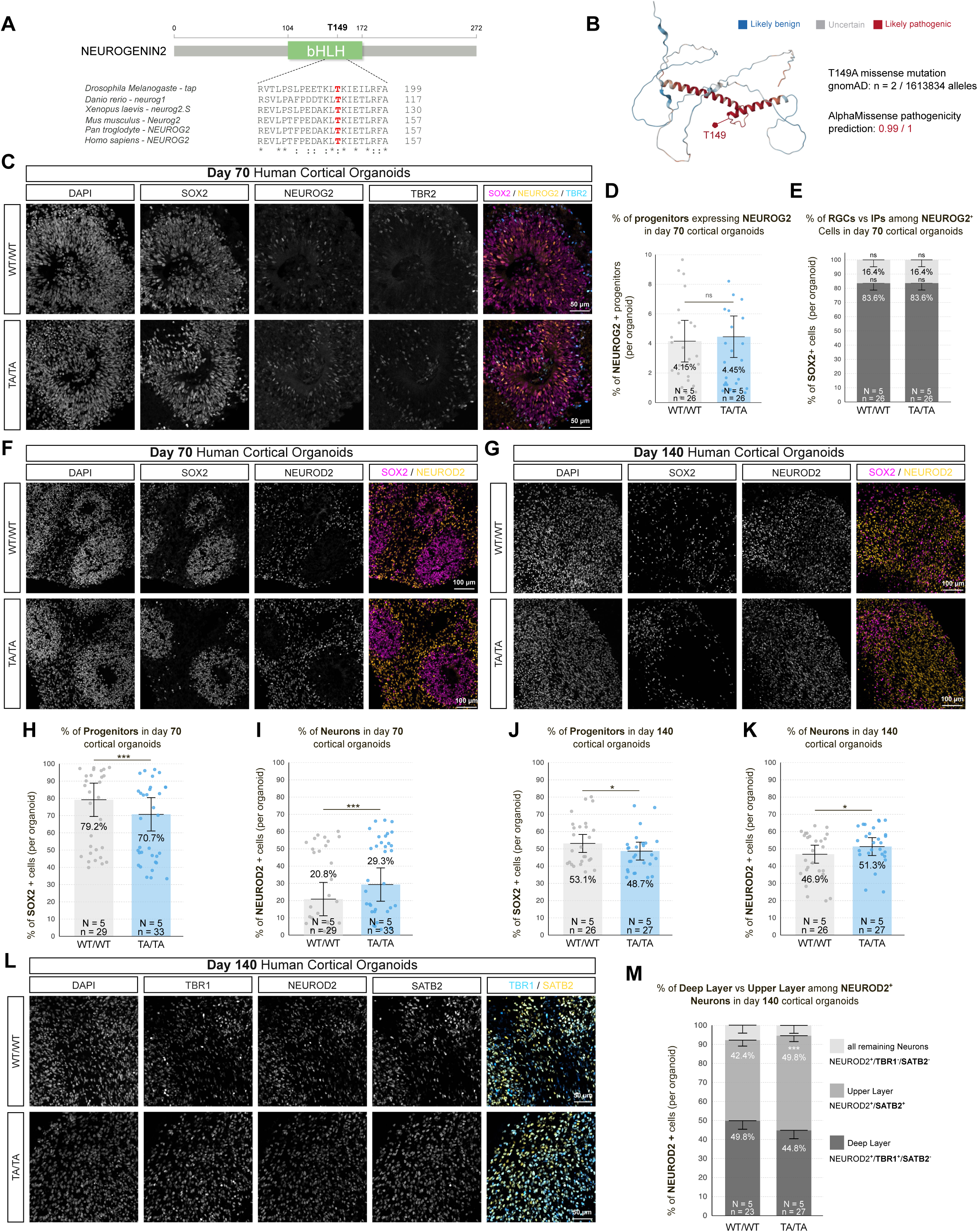
A conserved phosphorylation site regulates NEUROG2 proneural potency in human cortical organoids. **(A)** Multiple sequence alignment of the bHLH domain from NEUROG2 orthologs across species, highlighting the conserved threonine residue (T149 and equivalents) in red. Asterisks (*) denote fully conserved residues. Sequences shown correspond to: *tap* (Drosophila melanogaster), *neurog1* (Danio rerio), *neurog2.S* (Xenopus laevis), *Neurog2* (Mus musculus), *NEUROG2* (Pan troglodytes), and *NEUROG2* (Homo sapiens). **(B)** Lollipop plot of the human NEUROG2 protein (RefSeq NP_076924) showing the T149A missense variant (allele frequency 1.24×10⁻⁶), observed in two heterozygous individuals (gnomAD v4.0). The bHLH domain (residues 104–172) is shaded. AlphaFold2 predicts high pathogenicity for missense variants in this domain (score: 0.99/1). **(C)** Representative confocal images of day 70 human cortical organoids stained for DAPI (gray), SOX2 (magenta), NEUROG2 (yellow), and TBR2 (cyan). WT/WT: wild-type; TA/TA: NEUROG2 T149A phosphomutant clone. Scale bars, 100 μm. **(D)** Quantification of the proportion of progenitor cells co-expressing NEUROG2 at day 70, expressed as a percentage of all antibody-stained cells. **(E)** Proportion of radial glial cells (SOX2⁺/TBR2⁻) and intermediate progenitors (SOX2⁺ or SOX2⁻ but TBR2⁺) among all NEUROG2⁺ cells in day 70 organoids. **(F–G)** Representative confocal images of day 70 **(F)** and day 140 **(G)** organoids stained for SOX2 (magenta), NEUROD2 (yellow), and DAPI (gray). Scale bars, 100 μm. **(H)** Quantification of the proportion of SOX2⁺ progenitors at day 70, expressed as a percentage of all antibody-stained cells. **(I)** Quantification of the proportion of NEUROD2⁺ neurons at day 70, expressed as a percentage of all antibody-stained cells. **(J)** Quantification of the proportion of SOX2⁺ progenitors at day 140, expressed as a percentage of all antibody-stained cells. **(K)** Quantification of the proportion of NEUROD2⁺ neurons at day 140, expressed as a percentage of all antibody-stained cells. **(L)** Representative confocal images of day 140 organoids stained for TBR1 (blue), SATB2 (orange), and NEUROD2 (gray). Scale bars, 100 μm. **(M)** Proportion of deep-layer (TBR1⁺) and upper-layer (SATB2⁺) neurons among all NEUROD2⁺ cells in day 140 organoids.Bar graphs show adjusted means ± SEM derived from a linear mixed-effects model (group as fixed effect; differentiation batch as random effect). Pairwise comparisons were performed using estimated marginal means (emmeans); outliers (residuals > 2 SD) were excluded. N = number of independent differentiations; n = number of organoids. P < 0.05; p < 0.01; p < 0.001.

To test its importance, we used CRISPR/Cas9 genome editing to prevent T149 phosphorylation by replacing the T149 with an Alanine on both alleles (T149A, henceforth TA/TA) of our two iPSC lines, the WTSli008-A (Figure S4A) and GM25256*E (Figure S5A). For the main WTSli008-A iPSCs line, we confirmed with optical sequencing the absence of major DNA remodification post editing (Figure S4B & S4C). In mid stages hCOs, we found that the TA/TA mutation did not alter the proportion of NEUROG2+ cell (4.15% for the WT/WT and 4.45 % for the TA/TA) (Figure 3C & 3D), nor their distribution: with the majority of NEUROG2+ cells being RGCs in both WT/WT (83.6% ± 4.8%) and TA/TA (83.6% ± 4.8%) and the rest being IPs (WT/WT 16.4% ± 4.8%) and (TA/TA 16.4% ± 4.8%) (Figure 3e).

We next examined early (D70, Figure 3F) and late (D140, Figure 3G) neurogenesis. In D70 COs, we observed a significant 10% reduction in progenitors (SOX2+) with a corresponding increase in neurons (NEUROD2+) (Figure 3H & 3I), which was confirmed with our second iPSC line GM25256*E (Figure S5B-D). This significant change was persistent but milder at D140, with a 5% decrease in progenitors and a corresponding increase in neurons (Figure 3J & 3K). Finally, we assessed cortical laminar identities at D140 by staining for deep-layer (TBR1[) and upper-layer (SATB2[) markers (Figure 3L). Quantification revealed a significant shift in neuronal subtype specification: with WT/WT organoids exhibiting approximately equal proportions of deep-layer (49.8%) and upper-layer (42.4%) neurons, whereas TA/TA organoids showed a significant decrease in deep-layer neurons (44.8%) and a corresponding increase in upper-layer neurons (49.8%) (Figure 3M).

In summary, NEUROG2 T149A shows the reciprocal phenotypes to the NEUROG2 KO, increasing neuronal production throughout cortical neurogenesis, notably of upper-layer neurons.

### Inhibiting NEUROG2 Phosphorylation Drives RGCs into a Neurogenic State

So far, we have seen that NEUROG2 is mostly expressed by RGCs and that when the T149 is not phosphorylated, it drives increased neurogenesis in both mid and late stages of human cortical development. However, it remains unclear whether the increased neuronal output observed in the phospho-mutant arises from RGCs with direct neurogenesis or IPs with indirect neurogenesis. To address this, we performed single-cell lineage tracing using live imaging combined with correlative immunostaining ^47^. We infected D50–70 WT/WT and TA/TA cortical organoids with a GFP-expressing retrovirus and tracked individual progenitors (n = 386) over 48–96 hours, allowing us to capture 1-2 divisions per progenitor (Figure 4A). We then used immunostaining for cell fate markers (SOX2 for RGCs, TBR2 for IPs, and NEUROD2 for neurons) to determine the fate of GFP+ progeny, enabling us to classify each division as either proliferative or neurogenic (Figure 4A). RGCs were classified as undergoing proliferative divisions if both daughter cells remained RGCs or neurogenic divisions if at least one daughter became an IP or neuron (Figure 4B). Similarly, IP divisions were classified as proliferative (producing two IPs) or neurogenic (producing one or two neurons) (Figure 4C).

**Figure 4.**
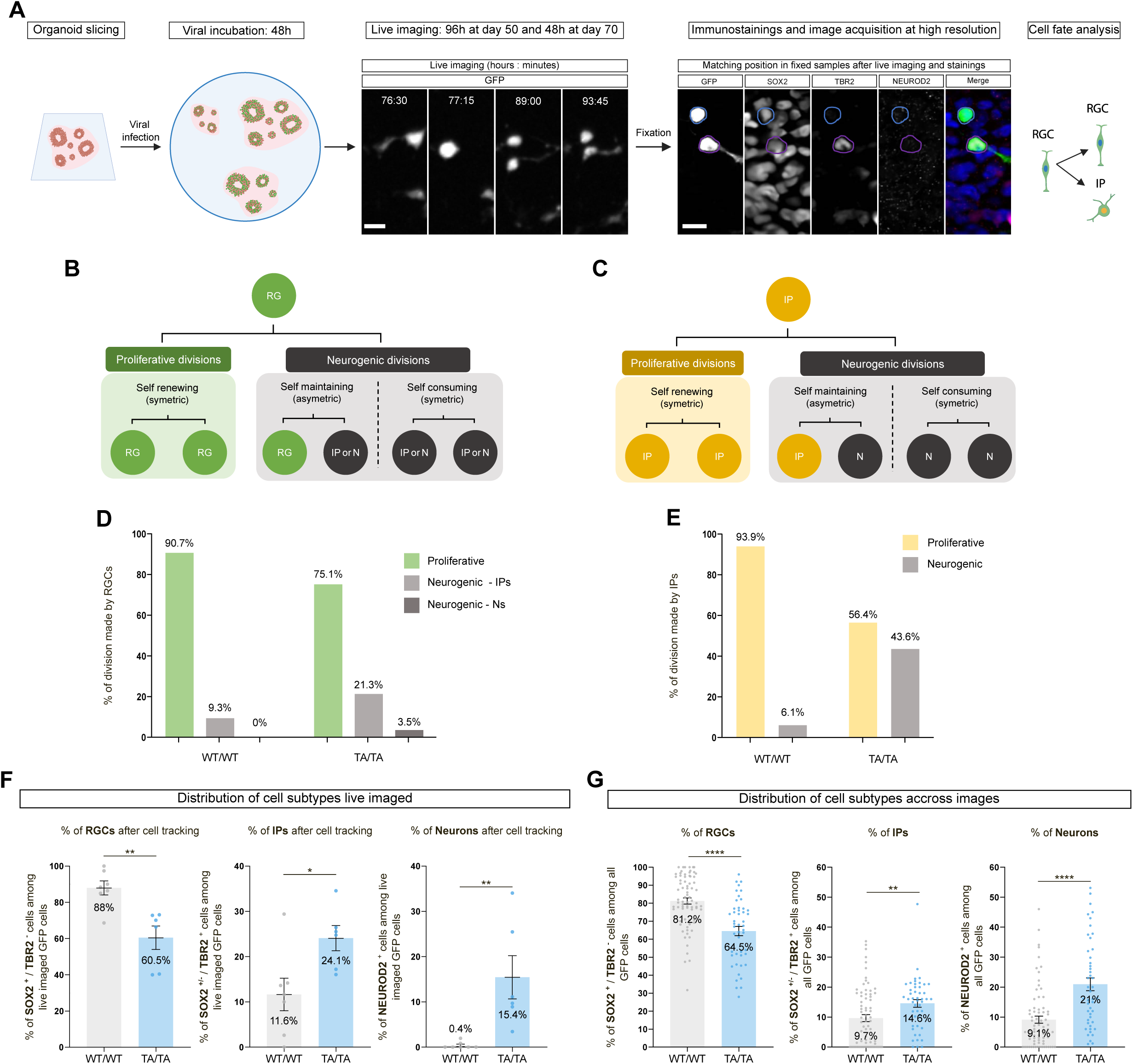
Phosphorylation of NEUROG2 at T149 regulates the balance between proliferative and neurogenic divisions in human cortical organoid progenitors. (**A**) Schematic overview of the correlative live imaging and immunostaining workflow. Human cortical organoids were sliced and infected with a GFP-expressing retrovirus for 48h at day 48 followed by live-imaging for 96 h. A second batch has been infected at day 50 or for 48 h followed by live imaging for 48 hours. After imaging, slices were fixed, stained, and re-imaged at high resolution. The division shown in the figure correspond to an asymetric neurogenic division. Data from day 50 and day 70 imaging were pooled. **(B)** Diagram summarizing possible division modes of RGCs, including proliferative (symmetric or asymmetric) and neurogenic divisions. **(C)** Diagram summarizing division modes of intermediate progenitors (IPs), which can self-renew or differentiate into neurons. **(D)** Quantification of division types for tracked WT/WT and TA/TA RGCs. **(E)** Quantification of division types for tracked WT/WT and TA/TA IPs. **(F)** Distribution of final cell subtypes (RGCs, IPs, Neurons) among all tracked progeny after live imaging. **(G)** Distribution of progenitor and neuronal subtypes among all GFP+ cells identified in fixed tissue sections, quantified across multiple fields of view from all infected organoid slices (N = 5 organoids per genotype). Bar plots represent means ± SEM. Pairwise comparisons performed using Student’s t-test. *p < 0.05; **p < 0.01; ***p < 0.001; ****p<0.0001.

In WT/WT RGCs, the vast majority of divisions (90.7%) were proliferative, with only 9.3% producing IPs and no direct neuronal production. In contrast, TA/TA phospho-mutant RGCs exhibited a marked decrease in proliferative divisions (75.1%) and a significant increase in neurogenic divisions, generating more IPs (21.3%) and neurons directly (3.5%) (Figure 4D), indicating enhanced direct and indirect neurogenesis. Tracing IP lineages revealed a similar trend. While most WT/WT IPs underwent proliferative divisions 93.9%, only 6.1% produced neurons. In TA/TA organoids, however, only 56.4% of IP divisions were proliferative, with 43.6% yielding neurons (Figure 4E). These results suggest that phospho-mutant RGCs not only produce more IPs but also bias these IPs toward neurogenic divisions, thereby amplifying neuronal output across successive generations.

This shift in division dynamics led to substantial changes in progenitor pools: live imaging revealed a reduction in RGCs from 88% to 60.5%, alongside increased proportions of IPs, from 11.6% to 24.1%, and neurons, from 0.4% to 15.4%, among TA/TA progenitor progeny over 2–4 days (Figure 4F). To validate these findings at a larger scale, we analyzed fixed organoid slices—including those not subjected to live imaging—by quantifying GFP+ cells across random fields of view. This analysis confirmed the same trend, with TA/TA organoids exhibiting decreased RGCs (from 81.2% to 64.5%) and increased IPs (from 9.7% to 14.6%) and neurons (from 9.1% to 21%) (Figure 4G). Together, these results demonstrate that blocking NEUROG2 phosphorylation shifts progenitors toward neurogenic trajectories, enhancing both direct and indirect neurogenesis.

### T149 attenuates NEUROG2’s proneural function

This raised the question of how the T149 residue regulates NEUROG2 activity, specifically, whether it alters NEUROG2’s intrinsic transcriptional capacity. To validate this, we first probed NEUROG2’s direct reprogramming potential with the rapid neuronal reprogramming protocol^48^ by using an all-in-one, rtTA-inducible lentiviral construct encoding the human NEUROG2 fused to eGFP that will reprogram iPSCs into neurons in less than a week. To prevent any positive feedback loop of NEUROG2, we used one of our NEUROG2 KO iPSCs clones and transduced it with two different lentiviruses, one expressing wild-type NEUROG2 and the other NEUROG2 T149A. We then selected the iPSCs that had been infected with lentiviruses through puromycin treatment to obtain stable clones with inducible NEUROG2 WT or NEUROG2 T149A. After NEUROG2 induction through doxycycline we quantified the fraction of MAP2[neurons among GFP[cells after six days in culture (Figure 5A). While WT NEUROG2 leads to a high conversion efficiency of 99.6%, the NEUROG2 T149A produced only 62.3% MAP2[neurons (Figure 5B), suggesting that, surprisingly, the T149A mutation decreases NEUROG2’s transactivation capacity. To test this directly, we compared the transactivation capacity of NEUROG2 WT and T149A in P19 cells, which lack endogenous NEUROG2 expression. We used luciferase reporters driven by either the Neurod1 promoter (long 2.2 kb variants; Figure 5C) or a multimerized E-box element from the Dll1 promoter (E2Z)[-luc (Figure 5D) as targets ^49,50^. Although NEUROG2 T149A retained the ability to activate both Neurod1 and Dll1 reporters, its activity was substantially reduced by ∼2-fold relative to wild-type NEUROG2, confirming the direct reprogramming results (Figure 5C-D & Figure S6A-F). Thus, paradoxically, preventing the phosphorylation of NEUROG2 T149 enhances its proneural function in cortical RGCs but it also substantially diminishes its transactivation potency.

**Figure 5.**
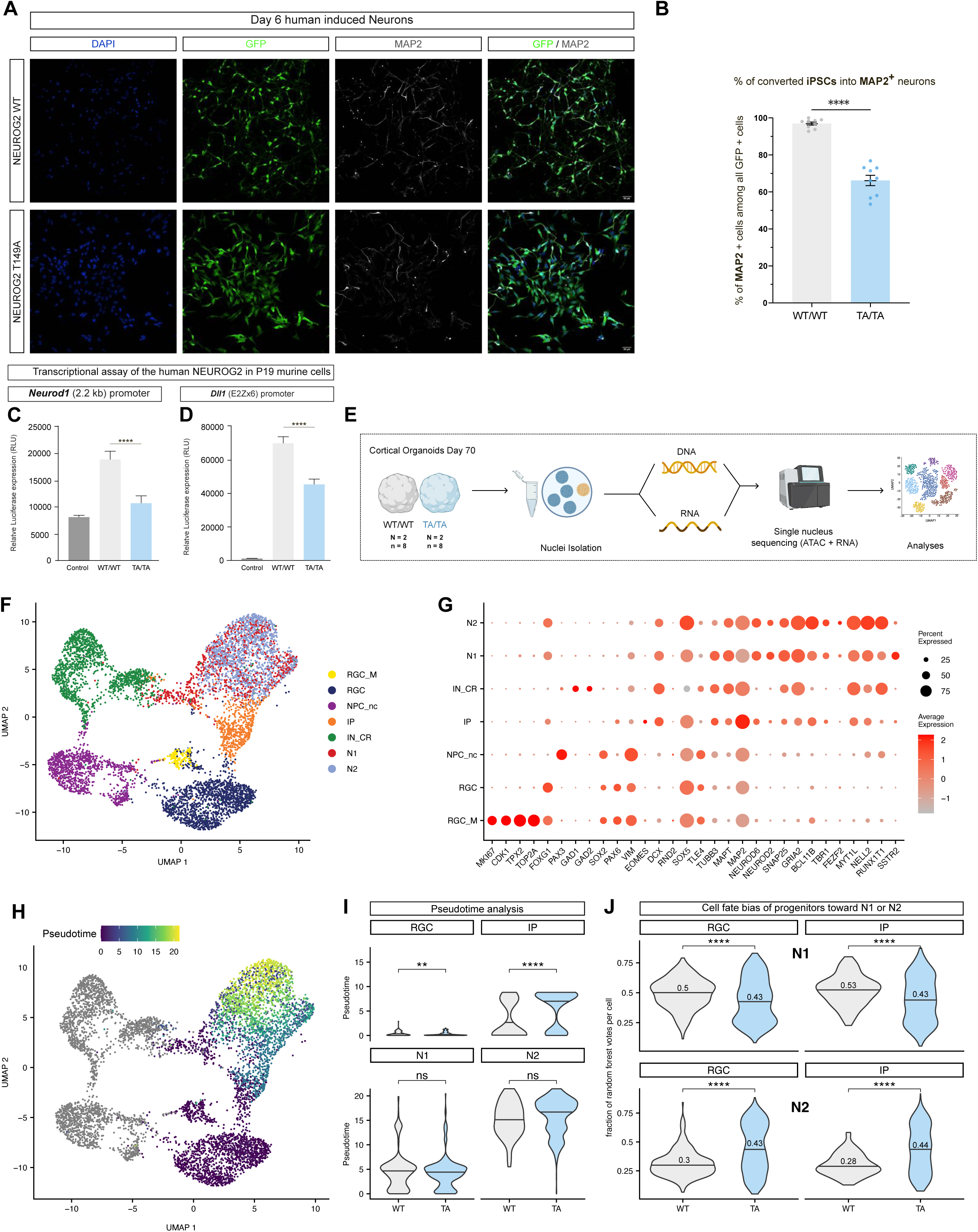
T149 regulate NEUROG2 activties. **(A)** Immunostaing for GFP (in green) and MAP2 (in gray) of converted iPSCs after 6 days in culture. **(B)** Quantification of MAP2 + neurons among all transfected GFP + cells. **(C)** Quantification of the relative expression of Luciferase in transcriptional assays for 1 representative experiment for the **(C)** Neurod1 2.2kb promoter and the **(D)** Dll1 promoter, in control: empty plasmid, WT: plasmid containing NEUROG2 WT, TA/ TA: plasmid containing NEUROG2 T149A. **(E)** Schematic of the single-nucleus multiomic experimental design. Day 70 WT/WT and TA/TA human cortical organoids (N = 2 differentiations per genotype, n = 8 organoids per differentiation) were processed for parallel snRNA-seq and snATAC-seq. Nuclei were isolated and analyzed using a joint modality embedding strategy. **(F)** UMAP embedding of WT/WT and TA/TA samples, showing integrated clustering of major cortical populations based on a weighted combination of RNA and ATAC profiles. Cell types include radial glia (RGC, RGC_M), intermediate progenitors (IP), early neurons (N1), late neurons (N2), and interneuron-like cells (IN_CR). **(G)** Scaled gene expression of canonical marker genes across identified clusters, shown as dot plots. Dot size represents the percentage of cells expressing the gene; color represents the average scaled expression. **(H)** UMAP embedding colored by pseudotime, with radial glia set as the root. Colors reflect the inferred developmental progression across cortical lineage states. **(I)** Pseudotime values across WT and TA conditions in RGCs, IPs, and neurons (N1 and N2). **(J)** Cell fate bias scores in RGCs and IPs toward either early (N1, top) or late (N2, bottom) neuronal states in WT and TA conditions. Values represent the median fraction of random forest classification votes supporting a given fate. **p < 0.01 ****p ≤ 0.0001, Wilcoxon test. Violin plots represent the median compared using Wilcoxon test and bar plots represent means ± SEM. *p < 0.05; **p < 0.01; ***p < 0.001; ****p<0.0001.

This paradox suggests that the increased neurogenesis observed in phosphomutant hCOs must result from another type of activity. More and more evidences suggest that NEUROG2 can induce a reshaping of the chromatin landscape by promoting DNA demethylation, increasing chromatin looping, and gene accessibility ^51–55^. Therefore, we examined gene accessibility and gene expression with paired single-nucleus multiome profiling (RNA-seq and ATAC-seq on the same nuclei, Figure 5E and Figure S7A-D) on eight control organoids and eight phospho-mutant organoids collected from two independent iPSCs 3D differentiation batches at day 70. We next integrated single-nucleus ATAC-seq and RNA-seq data into a unified UMAP projection, which resolved seven major cell populations (Figure 5F). Cortical clusters (FOXG1) contained the expected cell types, namely RGCs, mitotic RGCs (RGC_M: CDK1, TPX2, TOP2a), IPs, and two clusters of excitatory neurons at different stages of maturation (N1: SSTR2, NEUROD2, NEUROD6, and N2: MYT1L, NELL2, RUNX1T1, BCL11B) (Figure 5G).

To determine the effects of NEUROG2 T149A on the molecular program of cortical differentiation, we used Monocle3^56^ to reconstruct a pseudotemporal trajectory from the transcriptomes of all nuclei by designating RGCs as the origin of all the other cell subtypes (Figure 5H). The inferred path recapitulated the key stages of human cortical development starting with RGC, then IP, N1, and N2. Interestingly, TA/TA RGCs and IPs occupy significantly more advanced pseudotime positions than their WT/WT counterparts (Figure 5I, upper panel), indicating premature progression through progenitor states. Although N2 neurons exhibit higher pseudotime values than N1, consistent with their greater maturity, neither neuronal population showed a significant genotype difference (Figure 5I, lower panel).

These results suggest a priming of phospho-mutant RGCs towards more neurogenic states at the transcriptional level, but are they also biased towards faster maturation? To address this question, we applied FateID’s^57^ random-forest classifier to compute, for each RGC and IP, the probability of adopting an N1 versus an N2 identity. We observed that TA/TA RGCs and IPs are significantly more biased towards generating more mature neurons (N2) than the WT/WT cells (Figure 5J), confirming the more advanced neurogenic potential observed in cellular analyses.

### Phospho-mutant NEUROG2 remodels the RGC chromatin landscape

Taking advantage of the combination of single-nucleus RNAseq and ATAC-seq, we linked distal ATAC peaks, which are candidate cis-regulatory regions (cCREs), to their putative target genes using co-accessibility and correlated expression profiles (Figure 6A). Plotting the normalized accessibility of cCREs alongside the normalized expression of their linked genes for each cell subtype (RGC, IP, N1, N2) revealed both genes and cCREs with variable expression and accessibility patterns in TA/TA compared to WT/WT hCOs. To understand which transcription factors (TFs) might be underlying this variability, we performed TF-motif enrichment on linked cCREs (Figure 6B). We found an enrichment of motifs bound by different members of the JUN/FOS superfamily of transcription factors, collectively known as the AP-1 complex, in TA/TA RGCs compared to WT/WT RGCs (Figure 6B). Interestingly, the same AP-1 motifs were less enriched in TA/TA IP than in WT/WT IP cCRE, highlighting a transient and cell-specific chromatin remodeling. Consistent with this, JUN shows higher expression in RGCs than IPs (Figure S6E), and enhancers containing JUN motifs are associated with higher gene expression in RGCs while decreasing as RGCs enter neuronal differentiation (Figure S6F). Finally, regression-based enrichment analysis showed that while NEUROG2 is the most enriched motif in WT/WT RGCs, JUN, ATF4, and CTCF are among the most enriched TF motifs in TA/TA RGCs. This suggests a potential shift in the transcriptional program of RGCs as WT/WT RGCs rely more on NEUROG2 while TA/TA RGCs increasingly engage in AP-1-centered programs (Figure 6C & Figure S6G).

**Figure 6.**
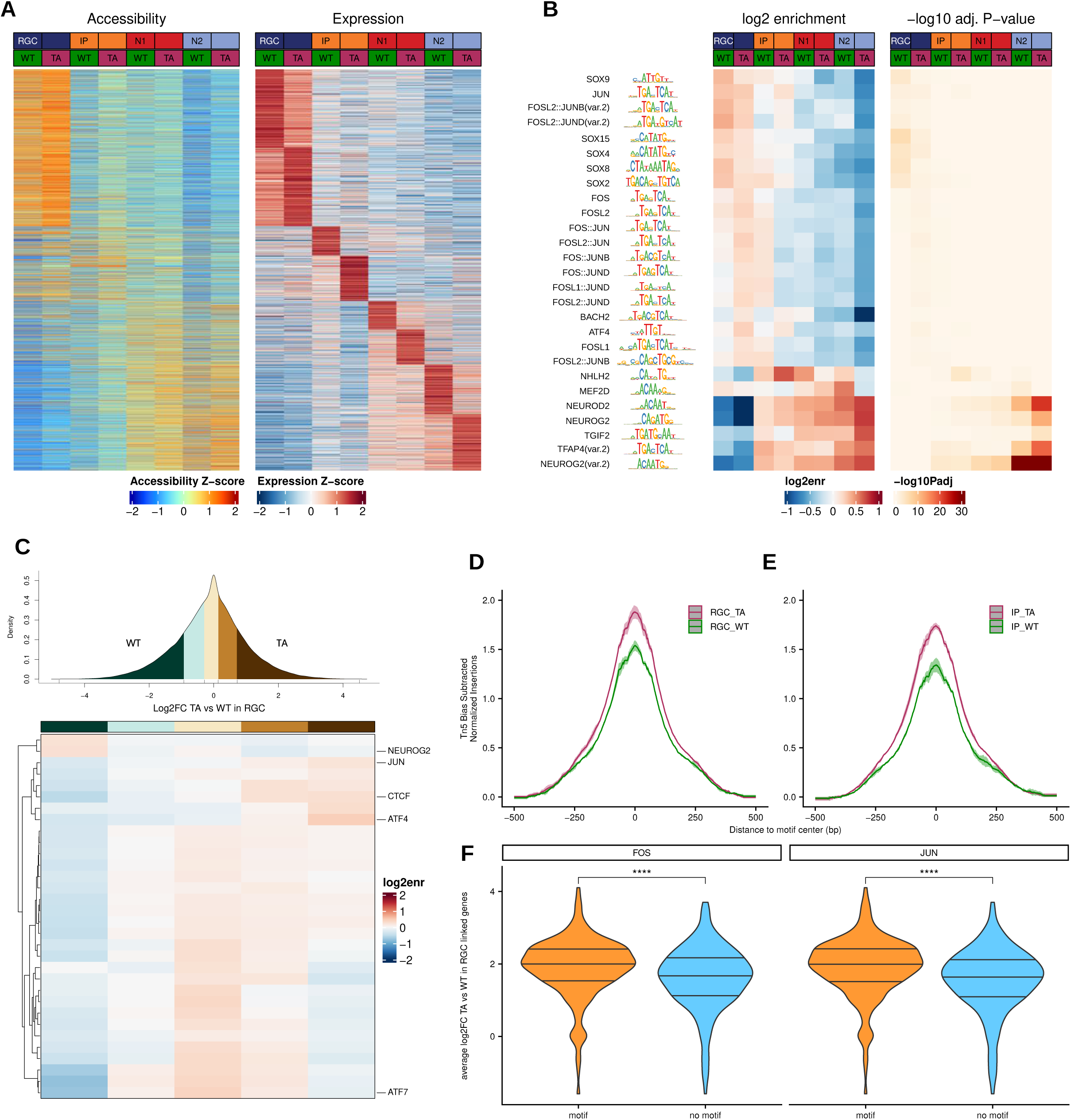
NEUROG2 T149 induces remodifications of the RGCs chromatin landscape in human cortical organoids. **(A)** Z-score heatmaps of chromatin accessibility (left) and gene expression (right) for enhancer–target gene pairs in radial glia (RGC), intermediate progenitors (IP), and neurons (N1, N2) in WT/WT and TA/TA conditions. Enhancers were linked to genes using co-accessibility and proximity criteria. **(B)** Transcription factor (TF) motif enrichment in distal regulatory elements linked to target genes from (A), per cell type and condition. Heatmaps show log2 enrichment (left) and –log10 adjusted p-values (right) for each motif. **(C)** Enrichment of motifs for expressed TFs in RGC marker peaks, binned by log2 fold change (TA/TA vs. WT/WT). Density plot (top) indicates the distribution of motif-containing peaks per bin; heatmap (bottom) shows motif enrichment (log2 enrichment) per bin, with key TFs labeled. **(D–E)** Aggregate Tn5 transposase insertion profiles (footprints) around JUN motif centers in RGC **(D)** and IP **(E)** marker peaks, comparing TA/TA and WT/WT conditions. **(F)** Violin plots showing pseudobulk average log2 fold change (TA/TA vs. WT/WT) in RGCs for genes linked to peaks containing (or lacking) FOS or JUN motifs. ****p ≤ 0.0001, Wilcoxon test.

Next, we assessed whether JUN family proteins were differentially bound to their motifs in TA/TA versus WT/WT progenitors using footprinting analysis, a method to detect physical binding of TF to their motifs. We found that the JUN footprint is stronger in TA/TA RGCs compared to WT/WT RGCs (Figure 6D), and while this footprint remains stronger in TA/TA IPs, it is reduced overall, and the JUN motif no longer appears among the enriched motifs in IP peaks (Figure 6E, Figure S6H). Finally, we quantified the effect of TF motif presence in cCREs on the expression of their linked target genes. For that purpose, RGC-specific genes were split into two sets. The first set comprised genes linked to distal enhancers having the AP-1 TFBS or one of its variants present in their sequence, and the second set comprised genes linked to enhancers without those motifs. Genes linked to RGC-specific peaks with FOS or JUN family motifs show significantly higher expression in TA/TA compared to those without these motifs (Figure 6F). This effect is specific to RGCs and is not observed in IPs or neurons (Figure S6F). Furthermore, this effect was observed only in RGCs but not in IPs or neurons (Figure S6F).

### NEUROG2 modulates the RGC gene regulatory network via JUN and FOS

Collectively, these observations establish that the phospho-mutant NEUROG2 leads to a remodeling of the chromatin landscape in RGCs, increasing accessibility and functional engagement of AP-1–associated regulatory elements. The enriched AP-1 motifs, JUN expression in RGCs, a stronger JUN footprint, and increased expression of AP-1 linked target genes suggest a potential rewiring of the gene regulatory network (GRN) of phosphomutant RGCs.

To explore the architecture of the GRN in sequenced RGCs, we used CellOracle^58^. The overall network topology remained similar between TA/TA and WT/WT RGCs, with a number of nodes, representing either a transcription factor or their target genes of 2025 nodes in TA/TA RGCs compared to 2005 nodes in WT/WT RGCs (Figure 7A, B). We also observed a slight increase in the number of edges, or regulatory connections, in the TA/TA RGCs with 9499 edges compared to 8993 edges in WT/WT RGCs (Figure 7A, c), suggesting a mild GRN expansion. To assess the influence of JUN and FOS on these networks, we computed three metrics: the number of direct connections a TF has (degree centrality), how often a TF acts as a regulatory bottleneck (betweenness centrality), and the number of a TF’s connections and the importance of its neighboring nodes (eigenvalue centrality). We found that both JUN and FOS showed increased centrality in TA/TA RGCs across the three parameters, indicating that these AP-1 factors regulate more target genes and are more central to key regulatory modules in the phospho-mutant context.

**Figure 7:**
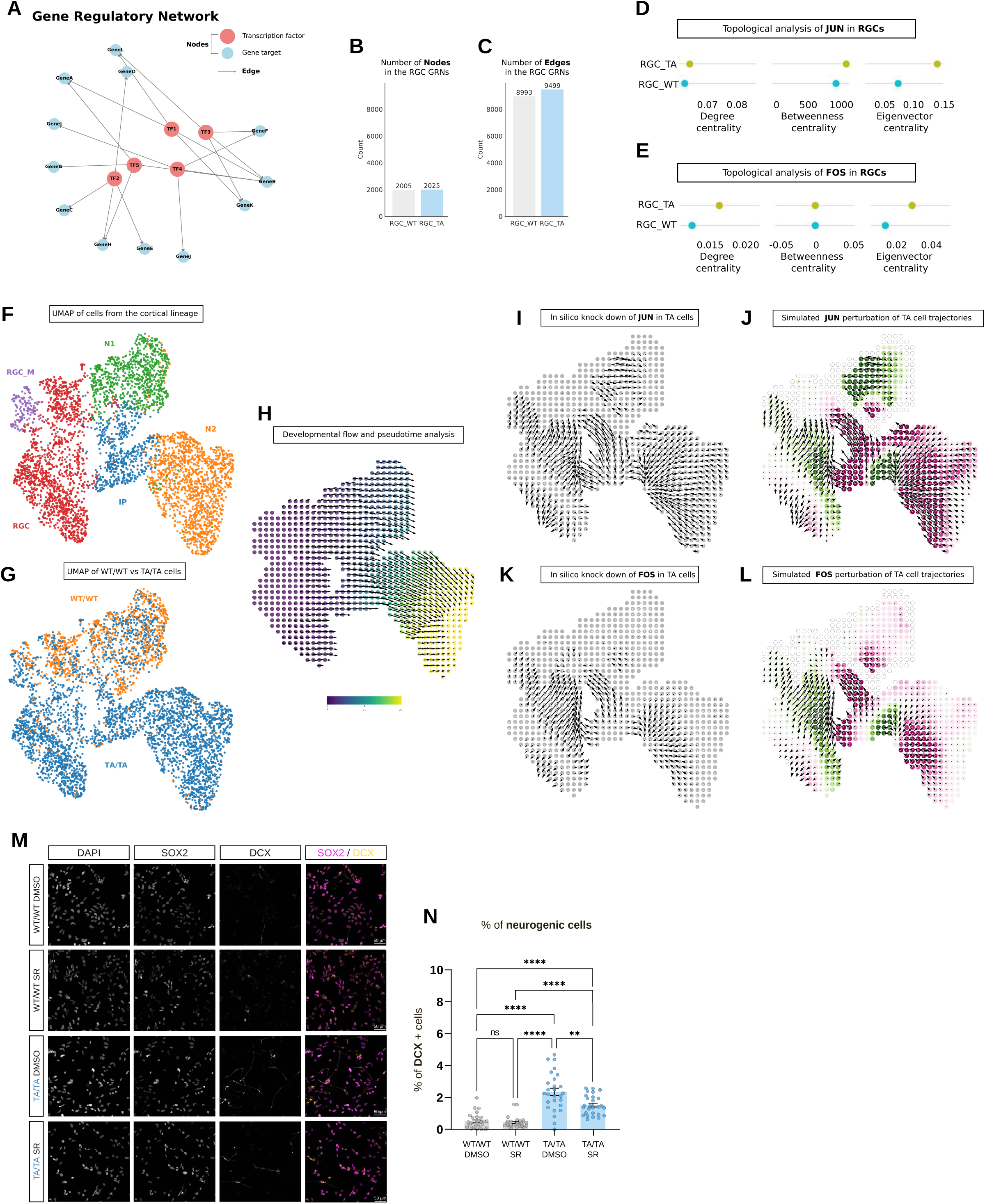
Gene regulatory network analysis identifies the AP-1 complex as a key regulator of neurogenesis in RGCs. **(A)** Schematic representation of a gene regulatory network (GRN), with nodes representing genes (TFs or targets) and edges representing regulatory interactions. **(B)** Bar plot showing the number of nodes in GRNs constructed from WT/WT versus TA/TA radial glial cells (RGCs). **(C)** Bar plot showing the number of edges in GRNs from WT/WT versus TA/TA RGCs. **(D)** Topological properties of JUN in GRNs from WT/WT and TA/TA RGCs: degree, betweenness, and eigenvector centrality. **(E)** Topological properties of FOS in GRNs from WT/WT and TA/TA RGCs, as in (D). **(F)** UMAP of all cortical lineage cells included in the multiomic analysis and used for GRN inference. **(G)** Same UMAP as in (F), colored by genotype (WT/WT or TA/TA). (H) Grid-based transformation of the UMAP embedding showing pseudotime overlay and CellOracle-derived 2D developmental vector field representing directionality of differentiation. **(I)** Grid plot showing the perturbation vector field following JUN knock-down in TA/TA cells. **(J)** Inner product of the JUN perturbation vectors with the developmental vectors across the grid space. Purple indicates inhibition of differentiation, green indicates promotion. **(K)** Grid plot showing the perturbation vector field following FOS knock-down in TA/TA cells. **(L)** Inner product of the FOS perturbation vectors with the developmental vectors, as in (J). **(M)** Representative confocal images of neural progenitor cells from cortical organoids treated with DMSO or the AP-1 inhibitor SR-11302, stained for SOX2 (magenta), DCX (yellow), and DAPI (gray). Scale bar, 25 µm. (**N)** Quantification of DCX+ cells shown in (M), expressed as a percentage of total cells. One Way ANOVA followed by Tukey-corrected pairwise tests: * p < 0.05; **p < 0.01; ***p < 0.001; ****p < 0.0001

However, it remains unclear whether AP1-centric transcriptional control drives the increased neurogenesis in phosphomutant RGCs in hCOs. To address this, we performed in silico knockdowns (KDs) of JUN and FOS to measure their potential impact on neuronal trajectories. We isolated cortical cells from our dataset and visualized their organization in UMAP. We found, that N1 and N2 neuronal populations form distinct clusters (Figure 7F) and WT/WT and TA/TA cells occupy different spaces (Figure 7G). To perform the perturbations, we built the developmental landscape using a grid point 2D vector field approach based on earlier pseudotime analysis (Figure 7H). Next, we performed in silico KDs of JUN and FOS (Figure 7I & 7k). By computing the inner product of the perturbation vectors with the developmental vectors of TA/TA cells, we found that JUN and FOS KDs reduce the differentiation potential of TA/TA RGCs and IPs (indicated by the purple color and back-flowing arrows) and revert them toward a WT/WT-like transcriptional state (indicated by the green color and upward flowing arrows) (Figure 7J & 7L).

To validate these analyses experimentally, we used 2D cultures of human RGCs kept under proliferative conditions. TA/TA RGCs displayed a ∼4-fold increase in the expression of the immature neuron marker DOUBLECORTIN (DCX) compared to WT/WT RGCs (Figure 7M & N), validating the increased neurogenesis of phosphomutant RGCs in 2D cultures. Inhibiting AP1 activity with the well-established molecule SR11302 ^59,60^ for 48 hours, corresponding to the length of one cell division in human RGCs, reduced the number of immature neurons (DCX+) by 1.8-fold (Figure 7N). Therefore, we demonstrated that NEUROG2 can promote neurogenic cell state transitions in human cortical cells through the AP-1 complex.

## DISCUSSION

One of the central questions in neurobiology is how species-specific features of the brain arise during development. While genetic innovations such as duplications of ARHGAP11B^18^ and NOTCH2NL^21,22^, or human-accelerated regions^27^, have been proposed to drive neocortical expansion, our findings suggest that post-translational modifications of conserved proteins like NEUROG2 may offer a complementary, reversible, mechanism for fine-tuning neurogenesis and thus potentially contribute to species-specific neocortical evolution.

### A robust and efficient Deep-Learning based image analysis workflow

To detect quantitative alterations with high confidence and at single-cell resolution, a central technical advance of our study was the development of a scalable image analysis pipeline, powered by deep learning, to quantify progenitor and neuronal dynamics across entire cortical organoid datasets. By combining automated nuclear segmentation with supervised machine learning–based classification, we analyzed over 100 million cells across thousands of sections. This unprecedented scale allowed us to detect subtle but robust shifts in progenitor fate and lineage progression that would be difficult to capture with traditional manual or low-throughput approaches. Beyond enabling our discoveries, this pipeline offers a powerful, adaptable resource for the general scientific community, allowing researchers to uncover hidden phenotypes in any model subjected to genetic perturbations, environmental challenges, or drug treatments.

### Species-Specific Roles and Post-Translational Proneural Modulation in Human Cortical Neurogenesis

At the cellular level, our findings reveal that NEUROG2 plays a broader role in human cortical neurogenesis than previously appreciated. In contrast to mice, where NEUROG2 is largely restricted to intermediate progenitors (IPs) and primarily promotes deep-layer neuron production, we show that in human models, NEUROG2 protein is predominantly expressed in radial glial cells (RGCs) and is required for the generation of both deep and upper-layer neurons. Importantly, this cannot be solely explained by a difference in the expression pattern, since electroporation of mouse Neurog2 into developing mouse RGCs accentuates deep-layer neurogenesis at the expense of upper-layer neurogenesis ^39^. This establishes NEUROG2 as a central regulator of human cortical neurogenesis and suggests a species divergence in its function. In addition, our findings support the hypothesis that decoupling protein activity from its expression could enable evolutionary changes in developmental timing, progenitor cell state transitions, and neuronal output. NEUROG2 phospho-mutant RGCs shift toward neurogenic divisions, both directly producing neurons and generating more IPs, which themselves are biased toward neurogenesis during mid-stages of cortical development. This effect persists into later developmental stages, specifically enhancing production of late born upper upper-layer neurons, a defining hallmark of human neocortical expansion. Importantly, this occurs without depleting the progenitor pool, suggesting that NEUROG2 phosphorylation status contributes to balancing self-renewal and differentiation in a way that maximizes neuronal output over extended periods.

### Uncoupling transcriptional and pioneer-like activities of a proneural factor

At the molecular level, Neurog2 has been known to function both as a transcriptional activator and as a pioneer-like factor capable of inducing reshaping of the chromatin landscape by promoting DNA demethylation, increasing chromatin looping, and accessibility ^51–55^. However, whether and how these dual activities, transcriptional activation and chromatin remodeling, are mechanistically separated and differentially regulated was unknown. We discovered that phospho-regulation of T149 uncouples NEUROG2’s transcriptional activity from its pioneer-like function to remodel the chromatin. Surprisingly, phospho-mutant NEUROG2 exhibited reduced transactivation capacity in direct reprogramming and reporter assays, despite promoting increased neuronal differentiation in cortical organoids. This apparent paradox is resolved by chromatin accessibility analyses, which revealed that inhibiting T149 phosphorylation leads to widespread premature opening of chromatin regions in RGCs, effectively priming progenitors for differentiation before they normally would. Importantly, this shift does not reflect accelerated maturation of neurons. Rather, it represents a forward temporal shift of progenitors into more differentiated trajectories, as demonstrated by pseudotime analysis. Furthermore, these chromatin changes show remarkable bias toward certain transcription factor networks, notably the AP-1 complex superfamily composed of different JUN and FOS. This was accompanied by increased accessibility and elevated expression of AP-1 target genes, as well as enhanced JUN transcription factor footprints in NEUROG2 phospho-mutant RGCs. Interestingly, these AP-1–associated chromatin changes were largely restricted to RGCs and faded in downstream IPs and neurons, suggesting that phospho-mutant NEUROG2 primes RGCs to enter neurogenic trajectories via selective enhancer activation. Pharmacological inhibition of AP-1 activity partially rescued the premature neurogenic phenotype after one cell division, confirming its direct functional role in driving the observed differentiation bias. Thus, it appears that NEUROG2 can drive neurogenesis through two different mechanisms. First, through its classical role by inducing the expression of genes such as NEUROD1, which promotes neuronal differentiation ^29^; and second, through increasing RGCs responsiveness to the AP-1 complex, a previously unknown role. We speculate that in humans these two roles maybe sequential, with the pioneering activity predominating in RGCs and the transactivation function required in IPs.

### The AP-1 complex: a hub for human cortical evolution

This mechanistic link positions AP-1 transcription factor networks as a central mediator of NEUROG2 phosphorylation effects. This aligns with accumulating evidence implicating AP-1 in species-specific aspects of cortical development. For instance, JUNB and FOSL2 have been shown to alter progenitor dynamics and extend neurogenesis in human models, while ectopic JUNB expression in mice confers human-like progenitor properties ^61^. Similarly, FOSL2 has been identified as a driver of human-specific gene regulatory networks in cortical progenitors ^62^. Thus, our results suggest NEUROG2 phosphorylation as an upstream modulator of a potential evolutionary hub: the AP-1 transcription factor network. By demonstrating that NEUROG2’s phosphorylation state selectively tunes AP-1–driven chromatin remodeling in human RGCs, our study suggests that NEUROG2 itself has been evolutionarily co-opted to engage AP-1 pathways in a human-specific context. Moreover, the evolutionary significance of AP-1 motifs extends beyond progenitors. Recent work has shown that species-specific changes in AP-1 binding sites underlie activity-dependent gene regulation in neurons ^63^, suggesting that AP-1 represents a generalizable axis of evolutionary plasticity throughout cortical development, from progenitor fate decisions to synaptic maturation. In this light, NEUROG2 phosphorylation emerges not merely as a developmental modulator but as a potential entry point into species-specific regulatory programs centered around the AP-1 complex.

### Context-specific regulation of proneural activity

Our study shows that phosphorylation of a single conserved residue, T149, can uncouple distinct activities of NEUROG2, thereby rewiring progenitor behavior and neuronal output in humans. This suggests that the interplay between kinases and phosphatases creates context-dependent windows of NEUROG2 activity that modulate neuronal production. Because phosphorylation depends on cellular energy and signaling states, this mechanism provides a direct link between metabolism, developmental cues, and progenitor fate. In this way, neurogenesis can be regulated at multiple levels by diverse key players, offering a tunable layer of control that may differ not only across progenitor subtypes but also between species. This raises a broader question: might evolution have harnessed reversible post-translational modifications to diversify cortical trajectories, increasing or limiting neuronal output in a species-specific manner? If so, the evolutionary expansion of the human cortex does not solely reflect the appearance of new genes, but also the flexible re-use of ancient proteins under novel regulatory dynamics—an elegant strategy for generating complexity without rewriting the genome. This conceptual framework offers a new lens on how evolutionary innovations in brain development can arise: through the convergence of genetic novelty with regulatory plasticity of conserved proteins.

## LEAD CONTACT

Further information and requests for resources and reagents should be directed to and will be fulfilled by the lead contact, Bassem Hassan.

## ACKNOWLEDGMENTS

We thank Dr. Nicolas Renier for comments and discussions. We thank the Paris Brain Institute ICV-iPSCs core facility for CRISPR/Cas 9 genome editing, the ICM-Quant facility for help with microscopy and especially David Akbar for his precious recommendations, and the Data Analysis Core for help with statistical analysis. This work was supported by Paris Brain Institute core funding, ANR grant NEURONPP (ANR-21-CE13-0041), the Roger De Spoelberch Prize (to BAH), the New Frontiers in Research Fund – Transformation, supported by the Canadian Tri-agencies – CIHR, NSERC, and SSHRC (to DSC), the Bettencourt Schueller Foundation Impulscience grant (to ADB), ERA-NET Neuron (MOSAIC), and European Research Council Consolidator Grant (EpiCortex, 101044469) (to BB). C.B.A. was funded by a Fondation pour la Recherche Médicale (FRM) post-doctoral fellowship.

## AUTHOR CONTRIBUTIONS

Conceptualization, J.P. and B.A.H.; methodology, J.P., B.A.H.,; Investigation, J.P.; T.D.; M.A.M.; L.R; M.V.S; N.D.; C.M.; C.B.A; H.M.; writing—original draft, J.P., T.D, B.B, B.A.H.; writing—review & editing, J.P. and D.C.; C.B.A; A.D.B; C.P.; B.B.; B.A.H.; funding acquisition, L.E.K; B.B; B.A.H.; resources, L.E.K; B.B. and B.A.H.; supervision, B.B. and B.A.H.

## DECLARATION OF INTERESTS

The authors declare no conflicting interests.

## METHOD DETAILS

### Experimental models

Two human induced pluripotent stem cell (iPSC) lines were used in this study: WTSIi008-A, obtained from the European Bank for Induced Pluripotent Stem Cells (EBISC), and GM25256*E, obtained from the Coriell Institute. Both lines underwent genomic characterization using Bionano optical mapping and were used for CRISPR/Cas9-mediated genome editing. WTSIi008-A served as the primary line for generating edited clones used in the main figures. Bionano analysis confirmed that no additional structural variants or recombination events occurred after CRISPR/Cas9 genome editing (Figure S3). The GM25256*E served as the secondary line to validate the results obtained from the WTSIi008-A in another genetic background.

Human fetal neocortical tissue at gestational week 18 (GW18) was obtained with prior informed consent and in full compliance with institutional and legal ethical guidelines. The protocol was approved by the French Biomedical Agency (Agence de la Biomédecine, approval number: PFS17-003). Fresh cortical tissue was collected from spontaneous miscarriages or medically indicated pregnancy terminations during autopsies performed at Robert Debré Hospital and Necker-Enfants Malades Hospital (Paris, France). A fragment of prefrontal cortex was dissected from one hemisphere and transported on ice to the laboratory. Samples were fixed in 4% paraformaldehyde (PFA) for 2 hours prior to further processing.

### iPSCs culturing and maintenance

iPSCs were cultured on Geltrex LDEV-Free hESC-qualified Reduced Growth Factor Basement Membrane Matrix (1%, Thermo Fisher Scientific, A1413302) coated dishes (B6 or B10 dishes) in mTeSR™ Plus (STEMCELL Technologies, #100-0276) supplemented with Antibiotic-Antimycotic (0,1%, Thermo Fisher Scientific, 15240062). The medium was changed every other day, and the iPSCs were passaged when iPSCs reached 80% confluency with 300µL of cGMP ReLeSR™ (STEMCELL Technologies, #100-0483) at various dilutions depending on the needs for the different experiments.

### CRISPR/Cas9 genome editing in human iPSCs

Guide RNAs (gRNAs) were designed using Benchling^64^. A total of 1×10[human iPSCs were nucleofected with a ribonucleoprotein (RNP) complex composed of 225 pmol each of crRNA and tracrRNA-ATTO+, and 120 pmol of Cas9 protein. Twenty-four hours after nucleofection, ATTO+ iPSCs were isolated by fluorescence-activated cell sorting (FACS) and seeded at low density (10–20 cells/cm²) on Synthemax II-coated surfaces in medium supplemented with CloneR2 to support clonal expansion. After one week, 192 individual iPSC clones were manually picked under a stereomicroscope and transferred to Geltrex-coated 96-well plates for expansion. Once confluent, clones were split for parallel cryopreservation and genomic DNA extraction. Genotyping was performed by PCR amplification followed by Sanger sequencing. List of primers and gRNA sequences used for the generation of the NEUROG2 KO clones of both iPSCs lines:

crRNA = GTCTGGTACACGATTGCAAA CGG

primer FWD and sequencing = CTGACCTTGGTTAGCACTGCC

primer REV = GGAATTGGAGGACACGGAGG

List of primers and gRNA sequences used for the generation of the NEUROG2 T149A clones of both iPSCs lines:

crRNA = AGCTCACCAAGATCGAGACC

primer FWD and sequencing = CTGACCTTGGTTAGCACTGCC

primer REV = CAGAAAGGCTACACCTGCCC

### Optical genome mapping

We used optical genome mapping (OGM) to evaluate genome stability in iPSCs following CRISPR-Cas9–mediated genome editing. This technique enables the detection of a wide range of chromosomal aberrations, both numerical and structural, with a resolution over 10,000 times greater than traditional karyotyping^65^. OGM has been shown to be a powerful tool for iPSC quality control^66^.

Briefly, ultra-high molecular weight DNA was extracted from frozen cell pellets using the Bionano Prep Blood & Cell Culture DNA Isolation Kit (Bionano Genomics, San Diego, CA, USA). DNA was then labeled with the Direct Label and Stain (DLS) enzyme using the Bionano Prep DNA Labeling Kit, following the manufacturer’s instructions. The labeled DNA molecules were loaded onto a Saphyr nanofluidic chip (Bionano Genomics) for linearization and imaged using the Saphyr instrument. The resulting images were converted into digital data and analyzed using Bionano Solve software.

*De novo* assembly was performed, followed by chromosomal analysis using two pipelines: a copy number variation (CNV) pipeline for detecting large, unbalanced events based on molecule coverage, and a structural variation (SV) pipeline for identifying both balanced and unbalanced rearrangements by comparing labeling patterns to a reference genome. Results were analyzed and visualized using Bionano Access software.

Variants detected in the edited lines (heterozygous and homozygous) but absent in the parental line were manually reviewed for validation.

### Generation of human cortical organoids from iPSCs

Cortical organoids were generated from human iPSCs using a previously reported protocol^42^ with modifications. hiPS cells were washed with PBS and incubated with 1mL of StemPro™ Accutase™ Cell Dissociation Reagent (Thermo Fisher Scientific, A1110501) for 5 min at 37 °C and dissociated into single cells. To obtain uniformly sized spheroids, approximately 3 × 10^6^ single cells were added per well in the AggreWell 800 plate (STEMCELL Technologies, 34815) with mTeSR™ Plus medium supplemented with Stemgent hES Cell Cloning & Recovery Supplement (1X, Ozyme, STE01-0014-500) and incubated at 37°C with 5% CO_2_. After 24 hours, spheroids from each microwell were collected by pipetting medium in the well up and down and transferred into Corning® non-treated culture dishes (Merck, CLS430591-500EA) in TeSR™-E6 (StemCell Technologies, #05946) supplemented with two inhibitors of the SMAD signaling pathway, dorsomorphin (5 μM, STEMCELL Technologies, #72102) and SB-431542 (10 μM, STEMCELL Technologies, #72234). From day 2 to day 5, TeSR™-E6 supplemented with dorsomorphin and SB-431542 was changed daily.

On day 6, the medium was replaced by Neurobasal™-A (Thermo Fisher Scientific, 10888022), B-27™ Supplement minus vitamin A (50X) (1X, Thermo Fisher Scientific 12587010), supplemented with GlutaMAX^TM^ (1%, Thermo Fisher Scientific, 35050038), 2-mercaptoethanol (0.1mM, Thermo Fisher Scientific, 31350010) and Antibiotic-Antimycotic (0.1%, Thermo Fisher Scientific, 15240062).

This medium was supplemented with 20 ng/mL Human Recombinant EGF, ACF (STEMCELL Technologies, #78136) and 20 ng/mL Human Recombinant bFGF, ACF (STEMCELL Technologies, #78134.1) and changed daily until day 12. From day 12 to day 24, the medium was changed every other day.

On day 25, the medium was replaced by Neurobasal^TM^ Plus (Thermo Fisher Scientific, A3582901), supplemented with B-27^TM^ Plus Supplement (50X) (1X, Thermo Fisher Scientific, A3582801), GlutaMAX^TM^ (1%, Thermo Fisher Scientific, 35050038), 2-mercaptoethanol (0,1mM, Thermo Fisher Scientific, 31350010), L-Ascorbic acid (200µM, Sigma-Aldrich, A4403), and Antibiotic-Antimycotic (0,1%, Thermo Fisher Scientific, 15240062). From day 25 to day 43, this medium was supplemented with Human Recombinant BDNF, ACF (STEMCELL Technologies, #78133), and Human/Mouse Recombinant NT-3 (STEMCELL Technologies, # 78074) and changed twice a week.

From day 43 until the last time points of interest, the same medium was changed twice a week without BDNF nor NT3.

### Cryopreservation of organoids

Organoids were collected in 2mL Eppendorf tubes, washed with PBS, and fixed in 4% PFA (Electron Microscopy Sciences, 15714) at RT for 6 hours in the dark. Organoids were washed several times in PBS before the incubation in sucrose 30% for at least 24h. When organoids are at the bottom of the tubes, they were collected and placed in the cryomolds for cryopreservation with Epredia™ Neg-50™ Frozen Section Medium (Fisher Scientific, 12688086) and frozen on dry ice. Organoids were then transferred to – 80°C upon slicing.

### Slicing of organoids

For immunostaining, COs were sectioned at a thickness of 14 µm using a Leica cryostat (Leica CM3050S) and collected on Superfrost Plus glass slides 25×75 (Fischer scientific, 11950657). The slides were stored at –80°C upon immunostaining.

### Stainings, image acquisition and analysis

To reduce the potential bias included by the experimenter and external conditions that vary throughout the years, all the cortical organoids of the same differentiation batch were stained, imaged and analyzed together. Here is the general pipeline from staining to image acquisition and analysis.

### Immunostainings

Slides were incubated at room temperature (RT) for 2 hours until completely dry. Slides were then washed with phosphate-buffered saline (PBS) to remove all cryoprotectant and blocked for 1 hour at RT with a solution containing 5% heat-inactivated horse serum (Thermo Fisher Scientific, 26050070), 3% bovine serum albumin (BSA) (Sigma-Aldrich, A9647-100G), and 0.3% Triton X-100 (Sigma Aldrich, X100-500mL). The sections were then incubated overnight at 4°C with primary antibodies diluted in the same blocking solution. After three washes with PBS for 5 minutes each, the sections were incubated with secondary antibodies and 4’,6-diamidino-2-phenylindole (DAPI) (1:5000, Sigma-Aldrich, D9564) diluted in the same blocking solution for 1 hour at RT. Following three additional washes with PBS, slides were mounted with VECTASHIELD Vibrance Antifade Mounting Medium (Eurobio Scientific, H170010) and stored at 4°C until imaging. Nail polish was added after polymerization of the mounting medium to ensure airtight sealing for long-term storage.

### Imaging of whole sliced organoids

Slides were positioned in the 4-slide plate (inverted-face down) and left at RT to warm before imaging with an inverted Nikon confocal AXR microscope. An initial low-resolution overview of the entire plate was acquired using a 4X objective (Resonant mode, 512×512 pixels, pinhole: 6 AU, high gain) in the 405 nm channel to capture DAPI signal. This step allows spatial registration of individual organoid slices and alignment with the identifiers assigned by the NIS software. High-resolution imaging was performed using a 20X air objective with Perfect Focus System enabled to maintain consistent z-plane positioning throughout acquisition. Each organoid slice was imaged as a tiled mosaic with a 15% overlap for better stitching using Galvano scanning (dwell time: 0.8 µs, pinhole: 1 AU, resolution: 2048×2048 pixels). Channels were acquired sequentially from longest to shortest wavelength (647, 555, 488, 405 nm) to prevent fluorophore cross-excitation. The mosaic dimensions were standardized across all slices based on the largest sample to ensure full coverage. Then image acquisition was launched after setting up all the parameters.

### Renaming of the images

After imaging, all slices were renamed following the same nomenclature throughout the entire project to store metadata in the title of the files. Here is the nomenclature:

Clone_Diff_Day_Staining_Objective_Parameters_OGnumber_SliceNumber

For example, a title would be: H9_dG_J70_SNd2_20Xa_g2dt08_OG7_s4 which corresponds to clone H9, a NEUROG2 T149A phospho-mutant from differentiation batch G fixed and stained at day 70 for the immunostaining of SOX2-NEUROD2 imaged with the 20X air objective using the mode galvano with an averaging of 2 and a dwell time of 0.8. This image corresponds to the slice 4 (s4) of organoid number 7 (OG7. The number of the organoid was set once for each batch and each time point and the number of slices corresponds to the number of slices obtained at the cryostat (usually 4 to 6 slices).

### Image processing

First, images were all opened manually in ImageJ for visual inspection. Any image with either an immunostaining too faint or with organoid slices not imaged entirely or the focus lost during acquisition were discarded from further analysis. Then the remaining images were manually cropped to save storage space and saved as tiff images for segmentation and object classification.

### Fine-tuning of StarDist, a deep learning algorithm for nuclear segmentation

#### Training

StarDist is a state-of-the-art machine learning-based method for cell segmentation that employs a star-convex polygon shape model to accurately segment nuclei^43,44^. While the pre-trained version of StarDist for fluorescent images yields very good results on images with sparse nuclei, it performs poorly on images with dense nuclei, as shown in Figure S2A. Thus, 134 images were taken using the Nikon AXR confocal microscope at different resolutions (Resonnant and Glavano mode) with either 1024×1024 pixels resolution or 2048×2048 pixels with the 20X air or 40X oil objectives to ensure the model to segment nuclei at different scale so it can be used on images taken from drosophila, mice and human samples. As StarDist internally uses convolutional neural network (CNN), the images don’t need to have the same x and y axis. Thus, images were cropped randomly to contain at least 100 to 700 cells each. All these images used for the training were manually annotated using Labkit, an ImageJ plugin. Furthermore, the original images and their corresponding masks used for training the model, which the Stardist authors published, were also downloaded. This resulted in a total of 447 images being downloaded from Stardist’s GitHub, corresponding to the stage1_train images from the Kaggle Data Science Bowl, which are available in full from the Broad Bioimage Benchmark Collection. The folder with the 447 images can be downloaded in a link provided in additional material. To enhance the training further, all the images were augmented before the training with techniques such as flipping the axis followed by rotations of 90°, 180° and 270° to multiply the number of images by 16. Then, in the training notebook, further augmentation was done by applying random intensity changes and Gaussian blur.

### Testing the model

In parallel, 9 images with various nuclei densities were taken as described above and manually labeled to evaluate the Stardist haug2 model fine-tuned. These images were not used for the training or the validation of our StarDist haug2 model. The model performances were assessed and are shown in Figure S2B. In short, the F1 score, which measures the model’s ability to avoid false alarms, i.e., predicting positive when the actual label is negative. The pretrained version of StarDist Versatile (fluorescent nuclei) scored an F1 score of 0.64, while the fine-tuned StarDist: stardit_haug2 has an F1 score of 0.83 (Figure S2C-E) on the test images. The model was then exported for use in Python but also in ImageJ or FIJI with version of TensorFlow 1.12 and 1.15, and in QuPath. The fine-tuned model, StarDist haug2 can be downloaded from the GitHub page of the lab and will be published on Bioimage.io for downloads. When used in Python, the Probability/Score Threshold and the overlap threshold are set automatically. In FIJI and QuPath, they have to be set manually to 0.51 and 0.3, respectively.

### Image segmentation

In Python, all images were then segmented using the fine-tuned model, StarDist haug2 on the DAPI channel (corresponding to channel 0 in the metadata of our images) with a tiling of 4×4 or 6×6 for faster processing of large images. The corresponding masks were saved either as uint16 when the image contains less than 65,535 nuclei or as uint32 bits when it contains more than 65,535 nuclei to optimize storage.

### Image selection for training the object classifier and assessing its performance

For better classification, the mean intensity of the nuclear markers of the different channels per image was calculated to plot the distribution of image intensities for a channel of interest across all the images. In general, 3 images were selected per genotypes for the training of the object classifier, with overall high, mean, and low intensities for the markers of interest.

### Object classification

We used a free and user-friendly software, ilastik, to train object classifiers ^45^. Training images and their corresponding masks were loaded in a new project for object classification. For training the random forest, we selected specific features among all the ones proposed by ilastik such as the mean intensity, the total intensity, the skewness of intensities, and the object’s area. Categories were created based on what type of cell population were stained. The training was done by assigning nuclei, and thus segmented objects, to each subcategory created in all the training images. The classifier measures all the previously selected features to establish a general profile of the cell. We then developed a simple python code to aggregate all the single files generated by ilastik for each image into a single file that corresponds to the RAW_DATA of the quantification done for the specific staining of the batch analyzed. With this file, we can then count the number of cells of each subcategory based on ilastik predictions per organoid slices and then the sum of all counted slice per organoid to get all the cells quantified per organoid. Organoids with less that 4 slices were discarded as the quantification might not reflect the full diversity of cell population across the entire organoid.

### Correlative immunostaining

The following technique is based on Coquand et al ^47^.

### Viral infection with retroviruses

Cortical organoids at both week 7 and week 9, corresponding to 55 and 63 days in culture, respectively, were collected and embedded in UltraPure low-gelling agarose (INVITROGEN, 16520-050) for subsequent slicing. A Leica VT1200S vibratome was used to slice the organoids into 250 µm – 300 µm thick sections at a 200µm/sec speed, while maintaining the samples in cold DMEM/F-12 (Thermo Fisher Scientific, 10565018). Slices were transferred in a 24 wells plate for retroviral transduction (MSCV-IRES-GFP, Addgene plasmid #20672) diluted at 1/30 in DMEM/F-12. After 2 hours of incubation, slices were washed using PBS and incubated in DMEM/F-12 supplemented with B-27 (Thermo Fisher Scientific, 10565018), N2 (1X, Thermo Fisher Scientific, 17502048), 10ng/mL FGF2 (STEMCELL Technologies, #78133.1), 10ng/mL EGF (STEMCELL Technologies, #78136), 5% fetal bovine serum (1X, Thermo Fisher Scientific, A5670801) and 5% horse serum (Thermo Fisher Scientific, 26050070) for 2 days before live imaging.

### Live Imaging

In order to follow RGCs and IP divisions over several days, slices were transferred onto a permeable membrane (Millicell Cell Culture Insert, 30 mm, hydrophilic PTFE, 0.4 µm, PICM0RG50, Merck) to create an air-liquid interface. The membrane was theninserted onto a 35mm glass bottom dishes (Fluorodish WPI, FD35-100) with 1mL of culture medium supplemented with EGF and FGF at a final concentration of 10 ng/mL under the membrane. Based on the GFP signal, positions were chosen and imaged every 15 minutes in 100µm stacks to capture migrating progenitors. However, the z step for each stack depends on the number of positions you have so that all the imaging fits within a 15-minute time window, where you have a good temporal resolution of cell divisions and migration to correlate the types of divisions with the daughter cell fate. At the end of the live imaging, low magnification images were taken with a 4X objective for alignment of the movies with the fixed, stained and mounted slices of hCOs.

### Immunostaining and movie processing

Subsequently, the slices were fixed using 4% PFA, with 1 mL applied under the membrane and 1 mL on top, to ensure maximal penetration of PFA. The slices were then washed with PBS for three 10-minute intervals and distributed into the wells of a 24-well plate. They were incubated for 1 hour in a blocking buffer consisting of PBS supplemented with 0.3% Triton-X and 2% donkey serum. Primary antibodies were incubated in the same solution overnight at 4°C on a shaking plate. Following this, the slices were washed using PBS + 0.05% Tween-20 before the incubation with secondary antibodies (diluted 1:1000 from our stock) in the same blocking buffer. The slices were then washed and mounted on Superfrost slides using Aquapolymount (18606-20, PolySciences). The slides were left in the dark at room temperature for polymerization and subsequently stored at 4°C until use.

### iPSCs infection with the NEUROG2 lentiviruses

The NEUROG2 KO clone 1 from the iPSCs line WTS8, lacking NEUROGENIN2 was infected with the two different lentiviruses containing either the WT version of the human Neurog2 or the NEUROG2 T149A phospho-mutant. On day 0, 60,000 iPSCs were incubated in a coated plate with 12 wells with the classical conditions as described earlier. On day 1, the cells were transduced with the lentiviruses at a MOI of 4 in mTeSR™ Plus (STEMCELL Technologies, #100-0276) supplemented with Stemgent hES Cell Cloning & Recovery Supplement (1X, Ozyme, STE01-0014-500) and 8µg/mL polybrene (Hexadimethrine bromide, Sigma-Aldrich, H9268-5G) for 3 days. After, we started the selection phase by changing the medium every day with increasing doses of puromycin (Sigma-Aldrich, P9620-10ML). For two days at 2µg/mL followed by three days at 1µg/mL. Then the selected iPSCs were then amplified and frozen in CryoStor CS10 (STEMCELL Technologies, 07930) until their use.

### iPSCs conversion into neurons

On day –1, 60,000 iPSCs were incubated in a well of a 12-well plate coated with Geltrex LDEV-Free hESC-qualified Reduced Growth Factor Basement Membrane Matrix (1%, Thermo Fisher Scientific, A1413302) in mTeSR™ Plus supplemented with hES. On day 0, the induction of NEUROG2 and NEUROG2 phospho-mutant expression was done by changing the medium with mTeSR™ Plus supplemented with 0.4µg/mL doxycycline (Sigma-Aldrich, D9891-5G). On day 1, the medium was replaced by Neurobasal Plus supplemented with BDNF and NT3 at a final concentration of 20 ng/mL with 0.4 µg/mL doxycycline. On day 3, the medium was replaced with the same medium as day 1, supplemented with cytosine β-D-arabinofuranoside (Sigma-Aldrich, C6645), to remove any potential dividing iPSCs that had escaped the selection process. After two days, on day 5, the medium was replaced with the exact same medium prepared on day 1. On day 6, neurons were fixed using PFA 4% for 10 minutes and washed three times very gently to prevent neuron detachment from the coverslips. Immunostaining using antibodies anti-GFP and anti-MAP2 was done to assess the conversion efficiency of each *NEUROG2* construct.

### Luciferase assay

P19 embryonic carcinoma cells were maintained in Dulbeccós modified Eagle’s high glucose medium (Sigma, D0422) supplemented with 10% Fetal Bovine Serum (Biowest, S008Y30304), 100 U/ml Penicillin-Streptomycin (Gibco, 15140122) and 2 mM L-Glutamine (Gibco, 25030149). Transfections were performed in quadruplicate in 48-well plates by using Lipofectamine 2000 (Thermo Fisher, 11668019) according to manufacturer’s protocol. Each well was seeded 1 day earlier with 6×104 cells and transfected with 200ng appropriate expression plasmid, 100ng luciferase reporter plasmid, and 200ng CMV-b-gal as internal control. Cells were lysed 24h after transfection with Glo Lysis Buffer (Promega, E2661), and extracts were assayed for luciferase and b-gal activities. Data are represented as means of quadruplicates.

### Quantification and statistical analysis

All statistical analyses of the data generated by our image analysis pipeline were quantified using RStudio with a linear mixed-effects model that accounts for the biological variability across independent iPSCs 3D differentiation batches. For each staining and timepoint, the percentage of neuronal or progenitor cells per organoid section was modeled using the lmer() function from the lme4 R package, specifying Group (e.g., genotype) as a fixed effect and Batch as a random intercept to correct for inter-experimental variation. Group-level comparisons were performed on the fitted models using the emmeans package, which computes estimated marginal means and pairwise contrasts via Students t-tests with Satterthwaite’s approximation for degrees of freedom. Model assumptions were assessed by visual inspection of residual plots. Outliers were defined as observations with standardized residuals exceeding ±2 standard deviations and were excluded before re-fitting the models.

All other statistical analyses were done with GraphPad Prism9, and normality of samples was verified by the Shapiro-Wilk test, followed by Student t tests when comparing only two conditions. Otherwise, we used a one-way ANOVA followed by a Tukey post hoc analysis for multiple comparisons.

For multiomic analyses, all statistical details, test and p-values, are given in the results sections and the figure legends.

### Multiomics analyses

#### QC and cell filtering

First, the raw sequencing data were demultiplexed with Cellranger (cellranger-arc-2.0.2) and mapped to the human hg38 reference genome (refdata-cellranger-arc-GRCh38-2020-A-2.0.0). On the feature-barcode matrix, QC metrics were computed in R using Seurat and Signac. Cells with minimally 2,000 and maximally 18,000 transcriptomic reads, minimally 2,000 and maximally 37,000 fragment counts, a minimal nucleosome signal of 0.2 and a maximum of 2, minimal TSS enrichment of 1 and maximal of 20, and minimally 1000 genes detected were retained (Fig S6 A-D). After QC filtering, there were 6,773 cells remaining. After peak calling with macs2^67^, non-standard chromosomes and blacklisted regions were removed.

### Dimensionality reduction and joint embedding

Next, the gene expression data were analyzed with Seurat. First, the transcriptomic counts were log-normalized, and the most variable features were selected. On the scaled and centered data, PCA was applied. The first 15 dimensions were used as input for a UMAP. Analogously, the chromatin accessibility data were analyzed with Signac and ArchR^68^. Normalization was done via term frequency-inverse document frequency (TF-IDF) normalization. The most variable peaks were identified, including features with >10 total counts. Those top peaks were scaled as well. Dimensionality reduction was done via singular value decomposition (SVD).

The first 15 and second to 15^th^ dimensions of the RNA and ATAC modality respectively were used to construct a weighted nearest neighbor (WNN) graph which was visualized via a UMAP representation. Doublet scores were computed with scDblFinder^69^. Harmony^70^ was used to correct for batch leading to an updated WNN and UMAP. Cells were clustered using Louvain. One of the clusters had a median doublet score of 0.99 and was therefore removed as putative doublet cluster. The final clustering comprises of 7 clusters, which were annotated using marker gene expression per cluster.

### Cell fate bias

The gene expression data of RGCs and IPs was tested per condition for a cell fate bias towards neurons (N1 and N2) with FateID^57^ using 5 cells per target cluster for classification per iteration and 10 cells per training set for classification

### Monocle 3 pseudotime

To obtain pseudotime values per cell type and condition, the monocle3^56^ algorithm was applied to the transcriptomic data using all cells in the RGC cluster as starting point.

### ATAC-seq analysis

Using ArchR, a reproducible peak set was established, and Transcription factors (TF) were annotated using JASPAR2020. Accessible peaks were linked to putative target genes (P2G) via Pearson correlation of peak accessibility and gene expression in a 500kb window. Only peaks that had a significant correlation (FDR < 0.1) and a minimal distance of 5kb to the TSS were retained as putative candidates of distal enhancers. Correlations were filtered for a value above 0.35 or below –0.35.

To decipher which TFBS in the peaks could be driving the gene expression in the target genes, a binned TF motif enrichment was done for the peaks linked to genes aggregated per cell type and condition with MonaLisa^71^.

### TF motif enrichment in RGC marker peaks

Marker peaks per cell type were identified using a Wilcoxon test using the TA condition as group and the WT condition as background while correcting for TSS enrichment and number of unique fragments per cell via ArchR. To identify TF binding sites, a TF motif enrichment in the RGC-specific marker peaks was done with MonaLisa using 5 equal sized bins based on the log2 fold change (LFC) of the marker peaks in the TA condition of RGC. Human PWMs were extracted from JASPAR2020 and filtered for expressed TFs. Additionally, a regression-based enrichment using randomized lasso stability selection was done for both RGC and IP, predicting the LFC of the Wilcox test of the marker peak selection with the TF motif hits in the peak sequences using a selection cutoff of 0.6. Motif footprints were determined in the marker peaks for JUN and FOS motifs using a modified version of the ArchR implementation. The footprints were normalized by subtracting Tn5 sequence bias.

### Effect of TF motif presence on gene expression

RGC specific genes and their linked peaks were extracted from the P2G matrix. TF motifs of the expressed TFs were added with Signac. Then, the peak set was split into a positive peak set containing at least one or more motifs of the TF of interest and a negative peak set containing none of these motifs. The RGC-specific genes were split into a positive and negative set depending on whether their linked peak contained these motif hits or not. These gene sets were compared across conditions in their average LFC across cells and in their expression per cell over the cell’s pseudotime values. To obtain LFC values, RGC TA vs WT was tested for differentially expressed genes in pseudobulk gene expression per cell type and condition with Seurat FindMarkers using TA as group and WT as background using DESeq2.

### Gene regulatory network construction

Gene regulatory networks (GRNs) were inferred using CellOracle. A base GRN was constructed from ATAC-seq fragments by performing motif scanning (JASPAR 2020) and associating transcription factors (TFs) with putative target genes based on promoter and enhancer accessibility. The multiomic dataset was first filtered to retain only cortical cells. Each cortical cluster identified earlier during the multiome analysis, was then subdivided by condition (WT or TA). GRNs were independently computed for each cell subtype and condition using CellOracle’s default thresholding parameters. To enrich the regulatory model, we manually extended the selected gene set to include the genes of the AP-1 complex and their putative target genes, resulting in a total of approximately 4500 genes used for GRN inference. No other specific filtering was applied.

### In silico gene perturbation

In silico perturbation analyses were performed using the GRNs derived from TA cells and plotted on the general UMAP computed after filtering for cortical cells. We simulated knockdowns of **JUN** and **FOS** by reducing their expression to 30% of baseline levels (0.3). CellOracle was used to compute the resulting vector fields representing predicted changes in gene expression and cell state across the existing low-dimensional space. These simulations were projected onto the precomputed pseudotime embedding to visualize how perturbation of AP-1 components would influence cell fate trajectories in the TA condition.

## ADDITIONAL RESOURCES

Downloading the images used for the pretrained versions of StarDist: https://github.com/stardist/stardist/releases/download/0.1.0/dsb2018.zip

Multiomic data can be found with GEO accession number: GSE305338

All the code for the image analysis pipeline can be found on GitHub at: https://github.com/Julien-Pgn/nuclei_quant

---

**Figure S1:**
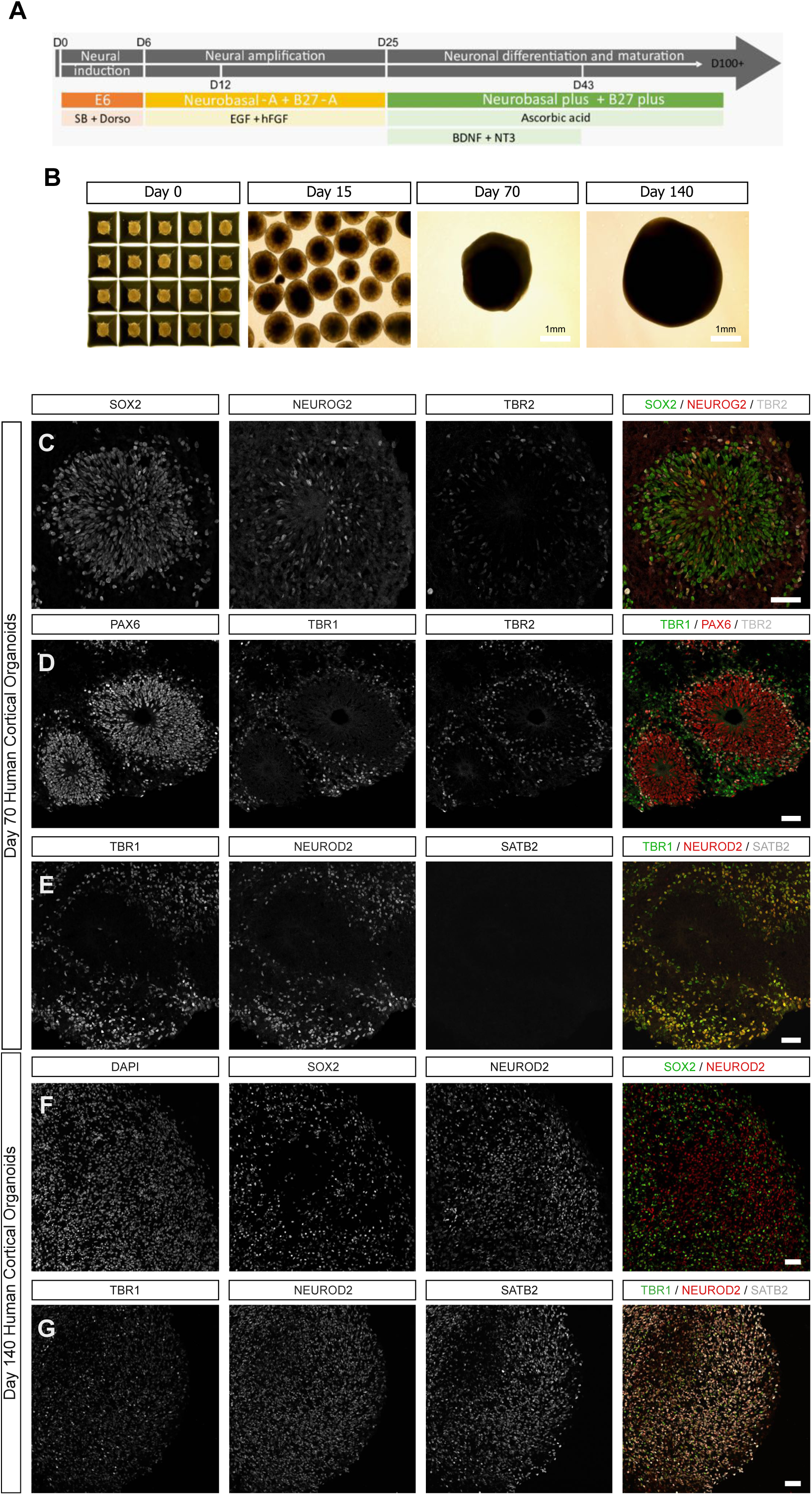
Validation of the 3D differentiation protocol of human iPSCs into cortical organoids. **(A)** Schematic representation of the protocol used, adapted from Sloan et al., 2018. **(B)** Brightfield images of human cortical organoids at different developmental stages: from day 0 in AggreWell plates to day 15 (rosette formation), day 70 (early/mid-stage), and day 140 (late-stage) of cortical organoid development. **(C)** Confocal images of hCOs at day 70 stained for SOX2, NEUROG2, and TBR2. **(D)** Confocal images of hCOs at day 70 stained for PAX6, TBR1, and TBR2. **(E)** Confocal images of hCOs at day 70 stained for TBR1, NEUROG2, and SATB2. **(F)** Confocal images of hCOs at day 140 stained for DAPI, SOX2, and NEUROD2. (G) Confocal images of hCOs at day 140 stained for TBR1, NEUROD2, and SATB2. In all images, scale bars represent 50 μm.

**Figure S2.**
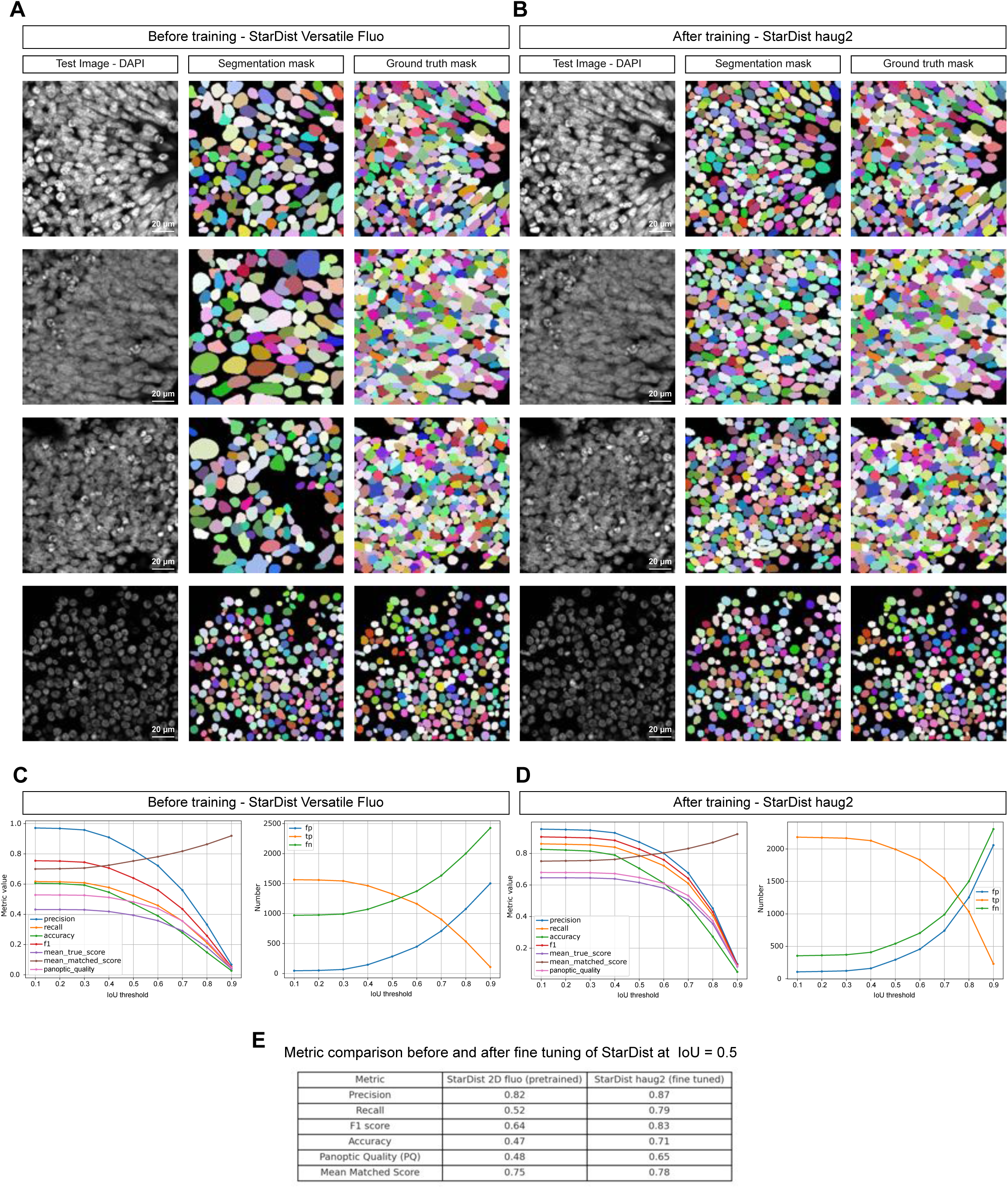
Fine-tuning of the StarDist model using manually annotated nuclei of human cortical organoids. **(A)** Representative test images (DAPI) used to evaluate the pretrained StarDist 2D Versatile Fluorescence model segmentation performance shown in the following panel with the annotated ground truth masks. Scale bars: 20µm. **(B)** Representative test images (DAPI) used to evaluate the fine tuned StarDist haug2 model segmentation performance shown in the following panel with the annotated ground truth masks. Scale bars: 20µm. **(C)** Performance metrics of the pretrained StarDist 2D model plotted across different intersection over union (IoU) thresholds. **(D)** Performance metrics of the fine-tuned StarDist haug2 model across IoU thresholds, showing improved prediction accuracy and precision. **(E)** Comparison of segmentation metrics at IoU = 0.5 for the pretrained and fine-tuned models, including Precision, Recall, F1 score, Accuracy, Panoptic Quality (PQ), and Mean Matched Score.

**Figure S3:**
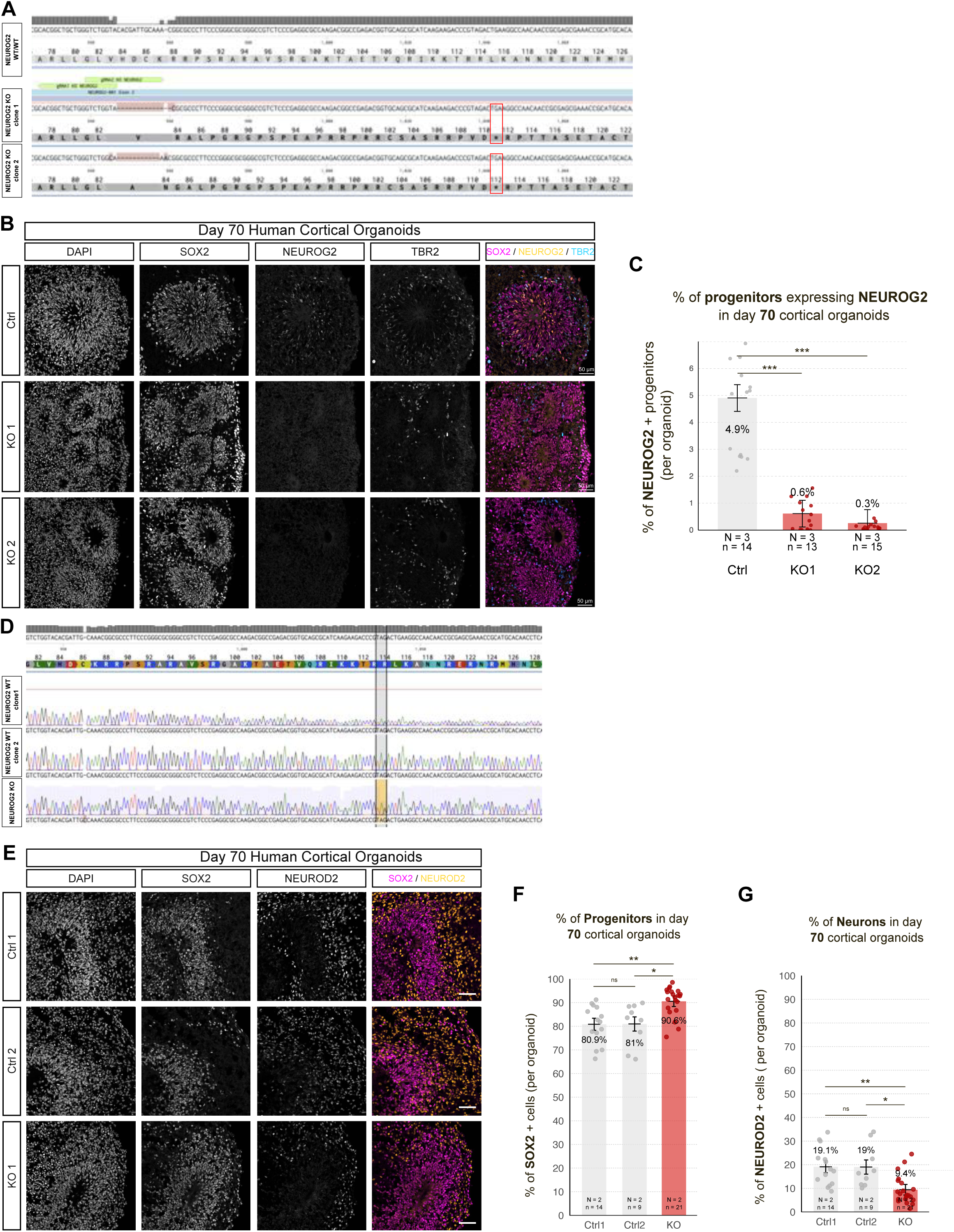
Validation of NEUROG2 CRISPR/Cas9 knockout and associated phenotype in two human iPSC lines. **(A)** Comparison of sequencing results from the control clone and two NEUROG2 KO clones of the WTSIi008-A iPSC line, showing deletions of 14 and 11 nucleotides, respectively, leading to premature stop codons (indicated by *). **(B)** Representative confocal images of human cortical organoids at day 70 stained for SOX2 (magenta), NEUROG2 (yellow), and TBR2 (cyan), plus DAPI (gray), for clones derived from the WTSIi008-A iPSC line: Ctrl: wild-type clone 1; KO1: NEUROG2 KO clone 1; KO2: NEUROG2 KO clone 2. Scale bars, 50 μm.**(C)** Quantification of the proportion of NEUROG2⁺ progenitors in day 70 cortical organoids, expressed as a percentage of all stained cells. **(D)** Comparison of sequencing results from two control clones and one NEUROG2 KO clone of the GM25256E iPSC line, showing a 1-nucleotide insertion leading to a premature stop codon (indicated by brackets). **(E)** Representative confocal images of day 70 human cortical organoids stained for SOX2 (magenta), NEUROD2 (yellow), and DAPI (gray), for clones derived from the GM25256*E iPSC line: Ctrl1: wild-type clone 1; Ctrl2: wild-type clone 2; KO1: NEUROG2 KO clone 1. Scale bars, 50 μm. **(F)** Quantification of the proportion of SOX2⁺ progenitors in day 70 cortical organoids, expressed as a percentage of all antibody stained cells. **(G)** Quantification of the proportion of NEUROD2⁺ neurons in day 70 cortical organoids, expressed as a percentage of all antibody stained cells.Bar graphs show adjusted means ± SEM derived from a linear mixed-effects model (group as fixed effect; differentiation batch as random effect). Pairwise comparisons were performed using estimated marginal means (emmeans); outliers (residuals > 2 SD) were excluded. N = number of independent differentiations; n = number of organoids. p < 0.05; p < 0.01; p < 0.001.

**Figure S4:**
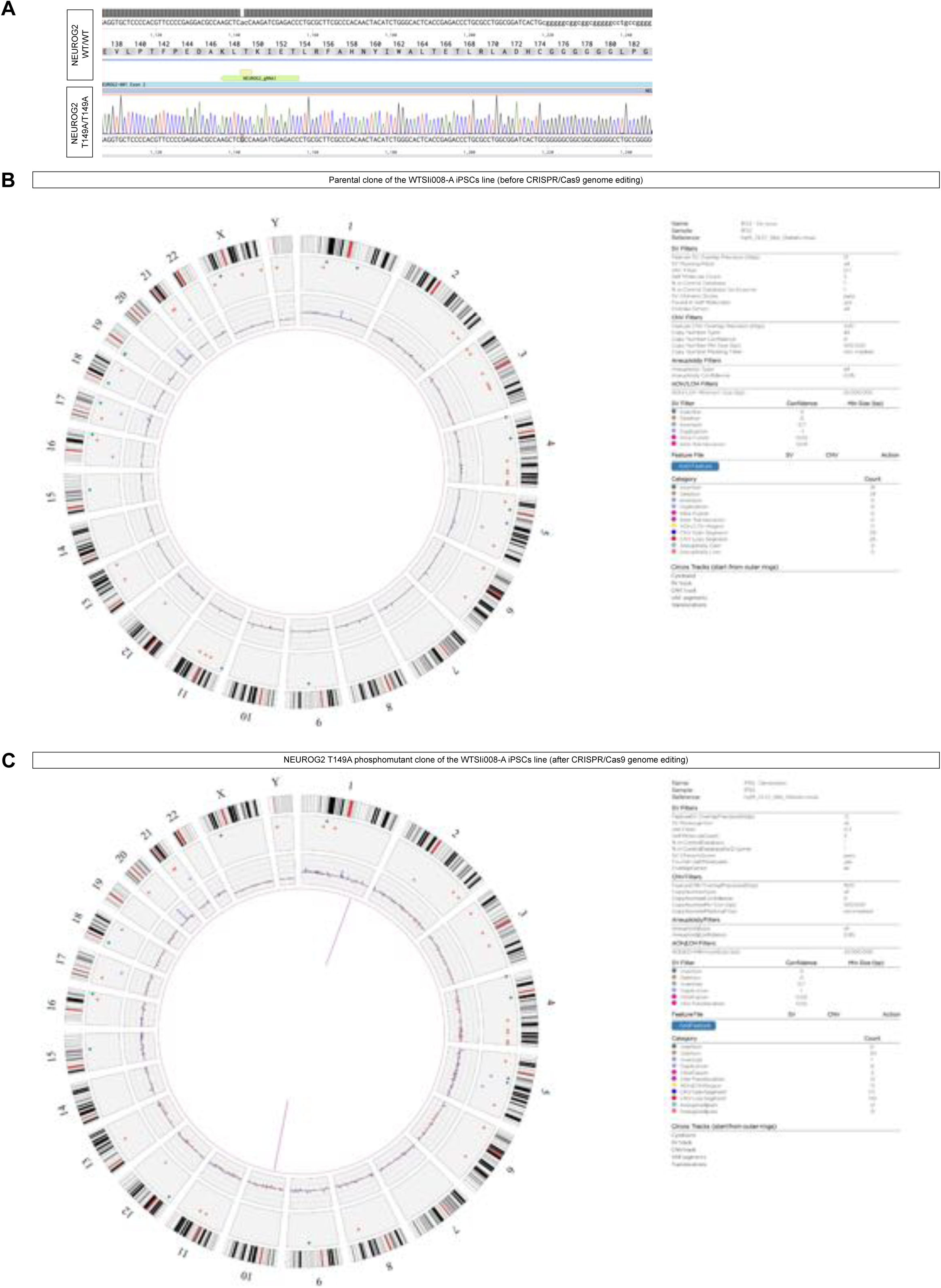
Validation of NEUROG2 T149A genome editing by CRISPR/Cas9 in the WTSIi008-A iPSC line. **(A)** Comparison of sequencing results from the NEUROG2 T149A phosphomutant and the wild-type NEUROG2 sequence after CRISPR/Cas9 genome editing. **(B)** Circos plot and associated filter table from optical genome mapping of the parental WTSIi008-A iPSC line using Bionano. Tracks are shown from outermost to innermost: chromosome ideogram, interstitial structural variants, copy number variants, and proximity lines for translocations, insertions, and inversions. In short a large duplication of the chromosome 20q has been found in the parental line. **(C)** Circos plot and corresponding filter table from genome analysis of the NEUROG2 T149A/T149A phosphomutant after CRISPR/Cas9 editing, validating the absence of off-target genomic alterations induced by the editing process. All minor detected structural variants compared to pannel (B) have been classified as false positive after careful manual verification.

**Figure S5:**
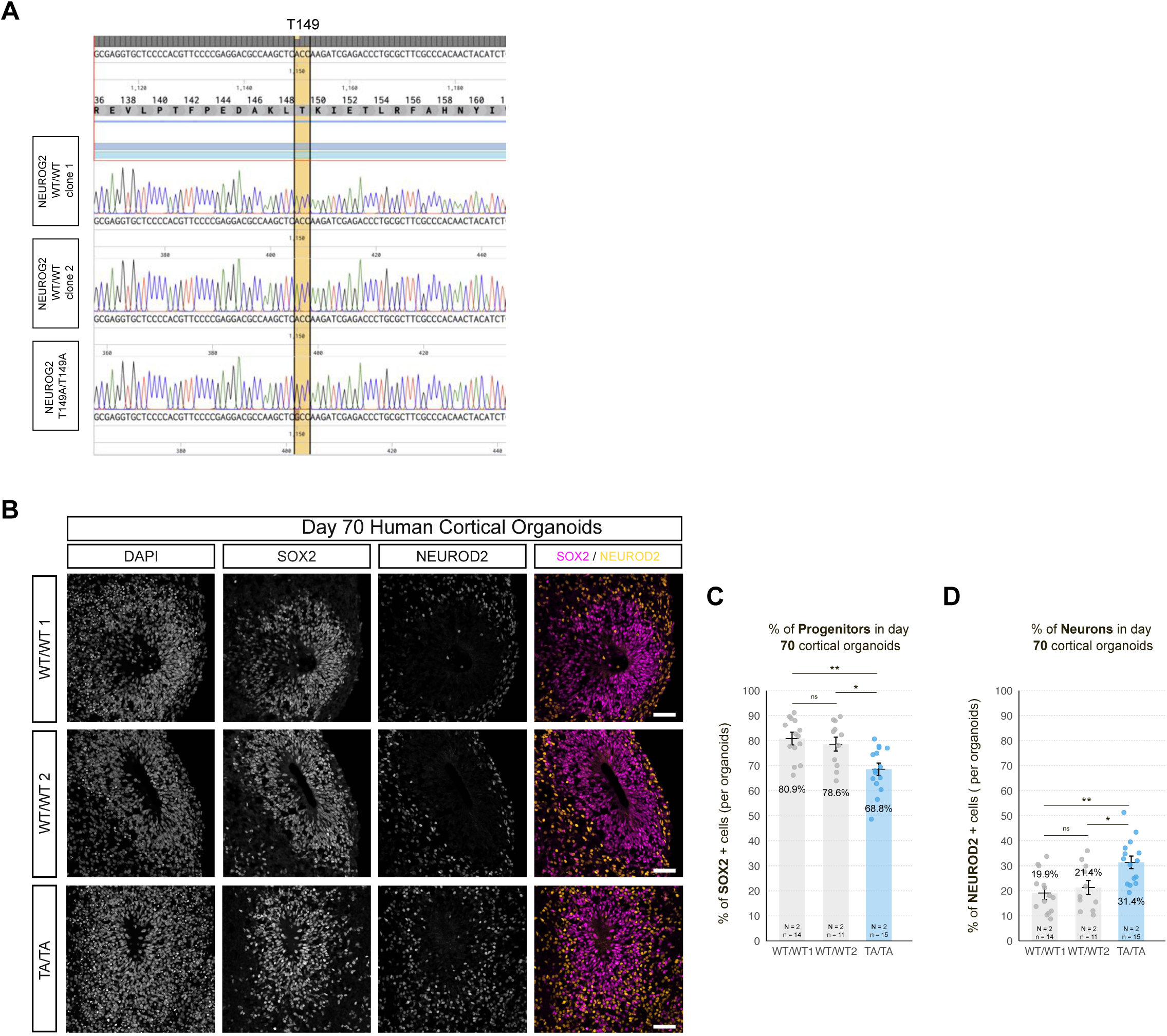
Validation of NEUROG2 T149A genome editing by CRISPR/Cas9 and associated phenotype in GM25256*E iPSC-derived cortical organoids. **(A)** Comparison of sequencing results from the NEUROG2 T149A phosphomutant (TA/TA) and two wild-type controls (WT/WT1 and WT/WT2), aligned to the reference NEUROG2 sequence. **(B)** Representative confocal images of day 70 human cortical organoids stained for SOX2 (magenta), NEUROD2 (yellow), and DAPI (gray), for clones derived from the GM25256*E iPSC line: WT/WT1: wild-type clone 1; WT/WT2: wild-type clone 2; TA/TA: NEUROG2 T149A phosphomutant. Scale bars, 50 μm. **(C)** Quantification of the proportion of SOX2⁺ progenitors at day 70, expressed as a percentage of all antibody-stained cells. **(D)** Quantification of the proportion of NEUROD2⁺ neurons at day 70, expressed as a percentage of all antibody-stained cells.Bar graphs show adjusted means ± SEM derived from a linear mixed-effects model (group as fixed effect; differentiation batch as random effect). Pairwise comparisons were performed using estimated marginal means (emmeans); outliers (residuals > 2 SD) were excluded. N = number of independent differentiations; n = number of organoids. p < 0.05; p < 0.01; p < 0.001.

**Figure S6.**
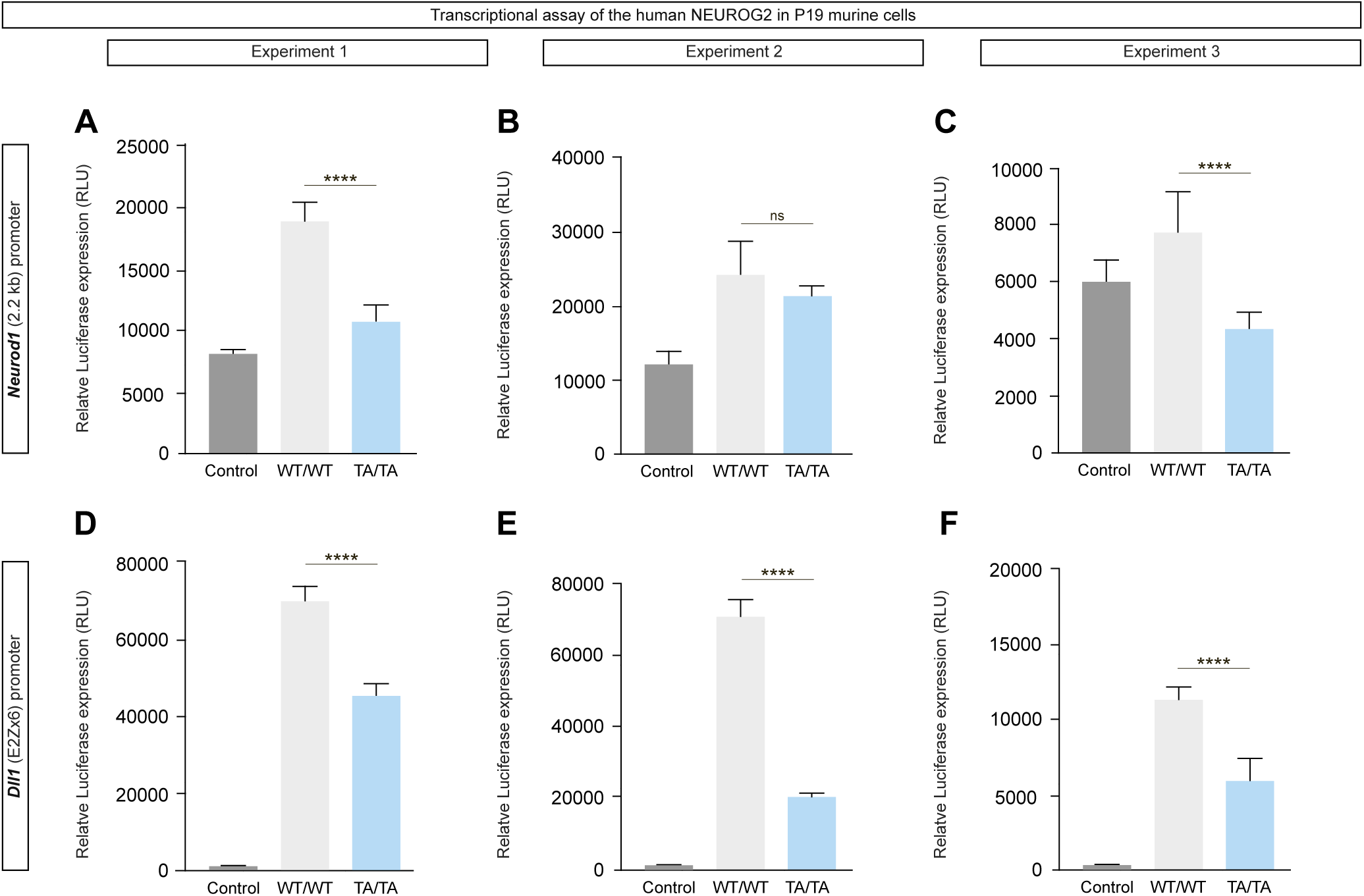
Unphosphorylated T149 induces a partial loss of transactivational properties of NEUROG2. (**A-C**) Quantification of the relative expression of Luciferase in transcriptional assays accross 3 independant experiment for the *Neurod1* 2.2kb promoter in control: empty plasmid, WT: plasmid containing *NEUROG2* WT, TA/TA: plasmid containing *NEUROG2* T149A. **(D-F)** Quantification of the relative expression of Luciferase in transcriptional assays accross 3 independant experiments for the *Dll1* promoter in control: empty plasmid, WT: plasmid containing *NEUROG2* WT, TA/TA: plasmid containing *NEUROG2* T149A. Bar plots represent means ± SEM. Pairwise comparisons performed using Student’s t-test. *p < 0.05; **p < 0.01; ***p < 0.001; ****p<0.0001.

**Figure S7.**
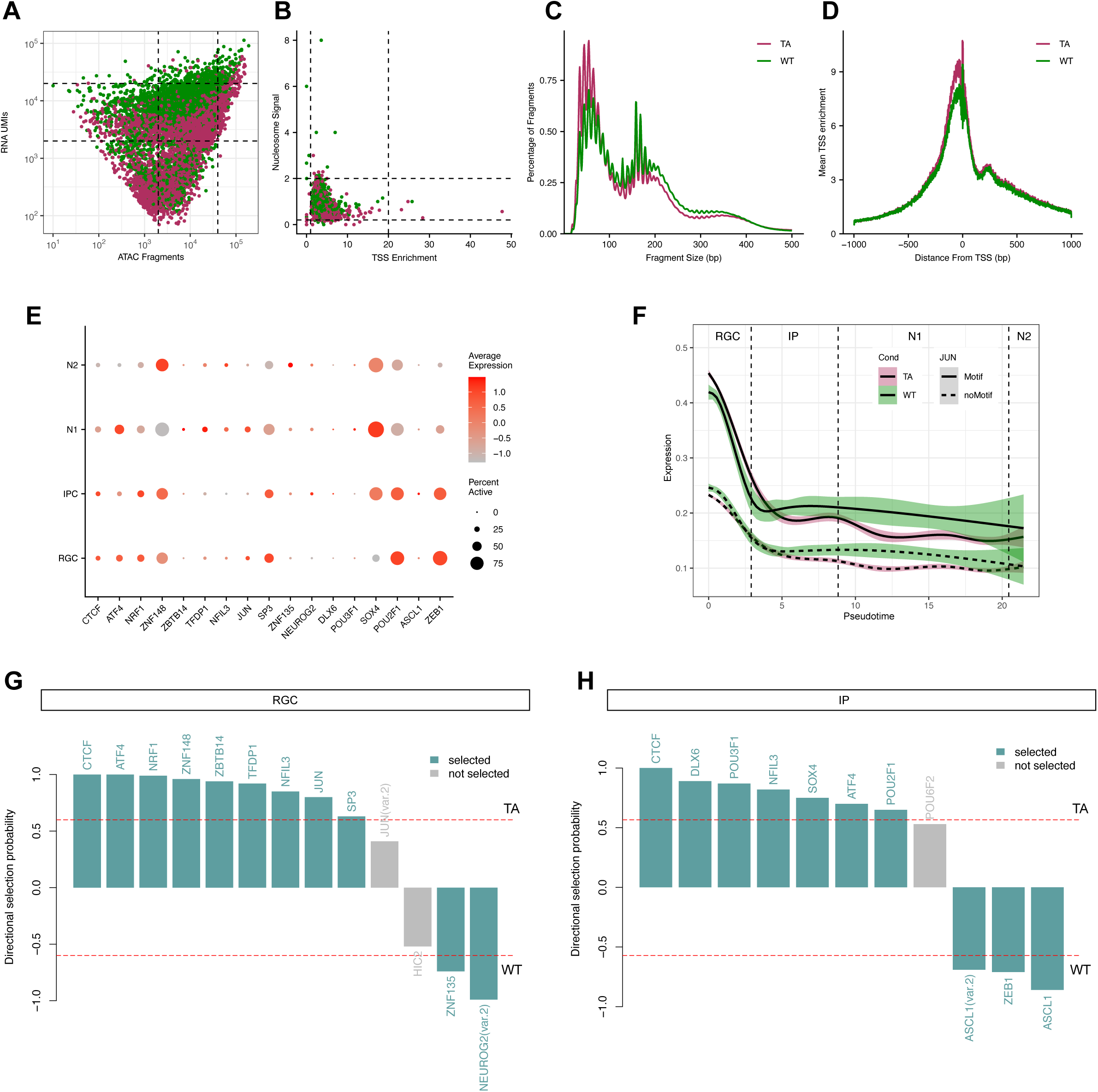
Quality control and transcription factor activity in single-nucleus multiomic data from human cortical organoids. **(A)** Scatter plot of RNA UMIs versus ATAC fragments per nucleus prior to filtering. Dashed lines indicate quality control thresholds used to exclude low-quality cells. **(B)** Scatter plot of nucleosome signal versus transcription start site (TSS) enrichment score before filtering. Dashed lines represent quality control cutoffs. **(C)** Fragment size distribution after filtering of low-quality cells, shown separately for WT/WT and TA/TA conditions. **(D)** Aggregate TSS enrichment profiles post-filtering, aligned to TSSs for WT/WT and TA/TA samples. **(E)** Dot plot showing scaled expression and percentage of cells expressing transcription factors with the highest selection probabilities from regression analysis in RGC and IP populations. **(F)** Gene expression dynamics of RGC-specific genes over pseudotime in IPs, stratified by presence (solid lines) or absence (dashed lines) of a JUN motif in the linked peak. TA/TA and WT/WT conditions are shown in pink and green, respectively. **(G–H)** Regression-based transcription factor motif enrichment analysis for expressed TFs, based on log₂ fold change (TA/TA vs. WT/WT) of marker peaks in RGCs **(G)** and IPs **(H)**. Bars represent directional selection probabilities: positive values indicate enrichment in TA/TA, negative values in WT/WT. TFs with significant selection are shown in color, with selected status indicated.

## Notes

### Competing Interest Statement

The authors have declared no competing interest.

### Summary of Updates

Correcting typos and minor textual changes

